# Structure of an insect gustatory receptor

**DOI:** 10.1101/2023.12.19.572336

**Authors:** Heather M. Frank, Sanket Walujkar, Richard M. Walsh, Willem J. Laursen, Douglas L. Theobald, Paul A. Garrity, Rachelle Gaudet

## Abstract

Gustatory Receptors (GRs) are critical for insect chemosensation and are potential targets for controlling pests and disease vectors. However, GR structures have not been experimentally determined. We present structures of *Bombyx mori* Gr9 (BmGr9), a fructose-gated cation channel, in agonist-free and fructose-bound states. BmGr9 forms a tetramer similar to distantly related insect Olfactory Receptors (ORs). Upon fructose binding, BmGr9’s ion channel gate opens through helix S7b movements. In contrast to ORs, BmGR9’s ligand-binding pocket, shaped by a kinked helix S4 and a shorter extracellular S3-S4 loop, is larger and solvent accessible in both agonist-free and fructose-bound states. Also unlike ORs, fructose binding by BmGr9 involves helix S5 and a binding pocket lined with aromatic and polar residues. Structure-based sequence alignments reveal distinct patterns of ligand-binding pocket residue conservation in GR subfamilies associated with distinct ligand classes. These data provide insight into the molecular basis of GR ligand specificity and function.

## INTRODUCTION

Animals rely on multiple chemosensory systems to adjust their behavior and physiology in response to changing external conditions and internal states. These include olfactory systems to recognize volatile chemicals and gustatory systems to detect ingested (often water-soluble) chemicals as well as low volatility, surface-associated chemicals. Detection relies on the cell-specific expression of distinct members of large families of membrane-spanning chemoreceptors^1^. Given their central role in transduction, determining the three-dimensional structures of chemoreceptors is important to understand the molecular mechanisms, including ligand recognition and selectivity, that govern chemosensory system function.

Insects are major sources of terrestrial biomass and biodiversity with major impacts on ecology and human health^2,3^. For example, some insects provide vital ecosystem services like pollination^4^, while others are disease vectors^5^ or agricultural pests that threaten food security^6^. Insect chemoreceptors influence feeding, reproduction, and other vital behaviors^1,7^, and are often accessible to external interventions (e.g., environmental chemical exposure). Therefore, knowledge of insect chemoreceptor structures and mechanisms can influence the development and use of agents for conservation and control efforts.

Among arthropods, including insects, Gustatory Receptors (GRs) have emerged as major mediators of chemosensation^8^, with 60 GR genes in *Drosophila melanogaster*^8^, 72 GR genes in the dengue vector mosquito *Aedes aegypti*^9^, and 65 GR genes in the silkworm *Bombyx mori*^10^. Many GRs are expressed in gustatory neurons, where they regulate feeding^1^. For example, in *D. melanogaster*, eight GRs contribute to sweet compound detection^11^, while six commonly-expressed GRs act alongside more than 20 additional GRs to detect bitter compounds^12^. GRs can also act in olfactory neurons; for example, disease-spreading mosquitoes use GR-mediated detection of carbon dioxide (CO_2_) to help locate human hosts^13^. GRs are also used to sense internal stimuli. In *D. melanogaster*, Gr43a acts as an internal nutrient sensor in the brain, monitoring hemolymph fructose levels to control satiety^14^ and egg production^15^. Thus, GRs detect a diverse array of chemical stimuli to control insect behavior and physiology.

GRs mediate chemosensation by forming ligand-gated cation channels^16–18^. GRs belong to the seven-transmembrane ion channel (7TMIC) superfamily, a large group of eukaryotic proteins that share transmembrane topologies, but little sequence identity^19^. In addition to GRs, insects contain other 7TMICs, including the Olfactory Receptors (ORs)^8,19^. The ORs diverged from within the GR family >400 million years ago and now form another large family of chemoreceptors in many insects (e.g., *D. melanogaster* encodes 68 GRs and 60 ORs)^8^. Although they share limited amino acid identity with GRs (<15%), ORs retain the 7TMIC organization and function as ligand-gated cation channels^20,21^. However, ORs are primarily involved in olfaction rather than gustation^8^. In most insects, individual ORs assemble with a common OR coreceptor (ORCO) to form functional ion channels^22^. While GRs lack an analogous coreceptor, many GRs also function in a combinatorial fashion^23^. However, some GRs form functional homomeric ion channels—these include *D. melanogaster* Gr43a and its *Bombyx mori* ortholog (Bm)Gr9, which are selectively activated by fructose^17^. While GR function has been extensively studied in animals, BmGr9 is one of the few GRs so far shown to form functional channels with native-like properties in heterologous expression systems^17^. For this reason, we chose BmGr9 as an initial GR family member for structural determination.

Although no experimental structure of a GR is available, there are structures of the fig wasp (*Apocrypta bakeri*) (Ab)Orco^24^ and the jumping bristletail *Machilis hrabei* (Mh)Or5, a homomeric OR that does not require an ORCO and is activated by a broad panel of odorants^25^. While GRs are evolutionarily related to ORs, their sequences are highly divergent and the agonists they recognize are chemically distinct (from water-soluble for many GRs to hydrophobic for many ORs). The similarity of GR and OR protein folds, and the extent to which GR and OR family members exhibit unique structural features that support their distinct functions, are unknown. Also unknown is whether the conformational changes induced by ligand agonists to open the ion pore differ between GRs and ORs, and how those changes are transmitted from the ligand-binding pocket to the pore.

Here we present a structure of BmGr9 in both the closed agonist-free state and in an open state in the presence of D-fructose. BmGr9 forms a homotetramer with a quadrivial pore architecture, and ligand binding results in ligand-binding pocket contraction and replacement of hydrophobic by hydrophilic residues at the extracellular gate of the ion pore. A set of four ordered phospholipids are found to lie horizontally in the plane of the membrane, with their headgroups interacting with conserved polar groups on the pore-lining transmembrane helix S7b and penetrating into the ion pore through fenestrations between subunits. Finally, we generate a structure-based sequence alignment and uncover distinct patterns of ligand-binding pocket residue conservation in GR subfamilies associated with distinct ligand classes. Together, these findings provide a high-resolution view of a GR family member, identify specific functions for conserved residues in long-established GR sequence motifs, and provide a starting point for more comprehensive analyses of this large and divergent family of sensory receptors.

## RESULTS

### Overall structure of an insect gustatory receptor

To determine the structure of BmGr9, we produced and purified N-terminally TwinStrep-tagged BmGr9 using HEK293 cells. When expressed in *Xenopus* oocytes, this construct is activated by D-fructose with an EC_50_ of 22 mM, but not by D-glucose (Figures 1A and S1A-D), consistent with the published EC_50_ value and specificity of wildtype BmGr9^17^. We produced detergent-solubilized BmGr9 (Figure S1E) and used single-particle cryo-EM to obtain a 2.85-Å density map (Figures 1B-C, S2 and S3) which we used to build and refine a structural model of BmGr9 (Figure 1D-E). We also purified BmGr9 in the presence of D-fructose (Figure S1F) and obtained a 3.98-Å density map which we used to model the D-fructose-bound conformation (Figures S4 and S5). We first describe the structural features of BmGr9 using the higher-resolution agonist-free structure, then compare the D-fructose-bound and agonist-free structures to define the fructose-mediated structural changes.

**Figure 1.**
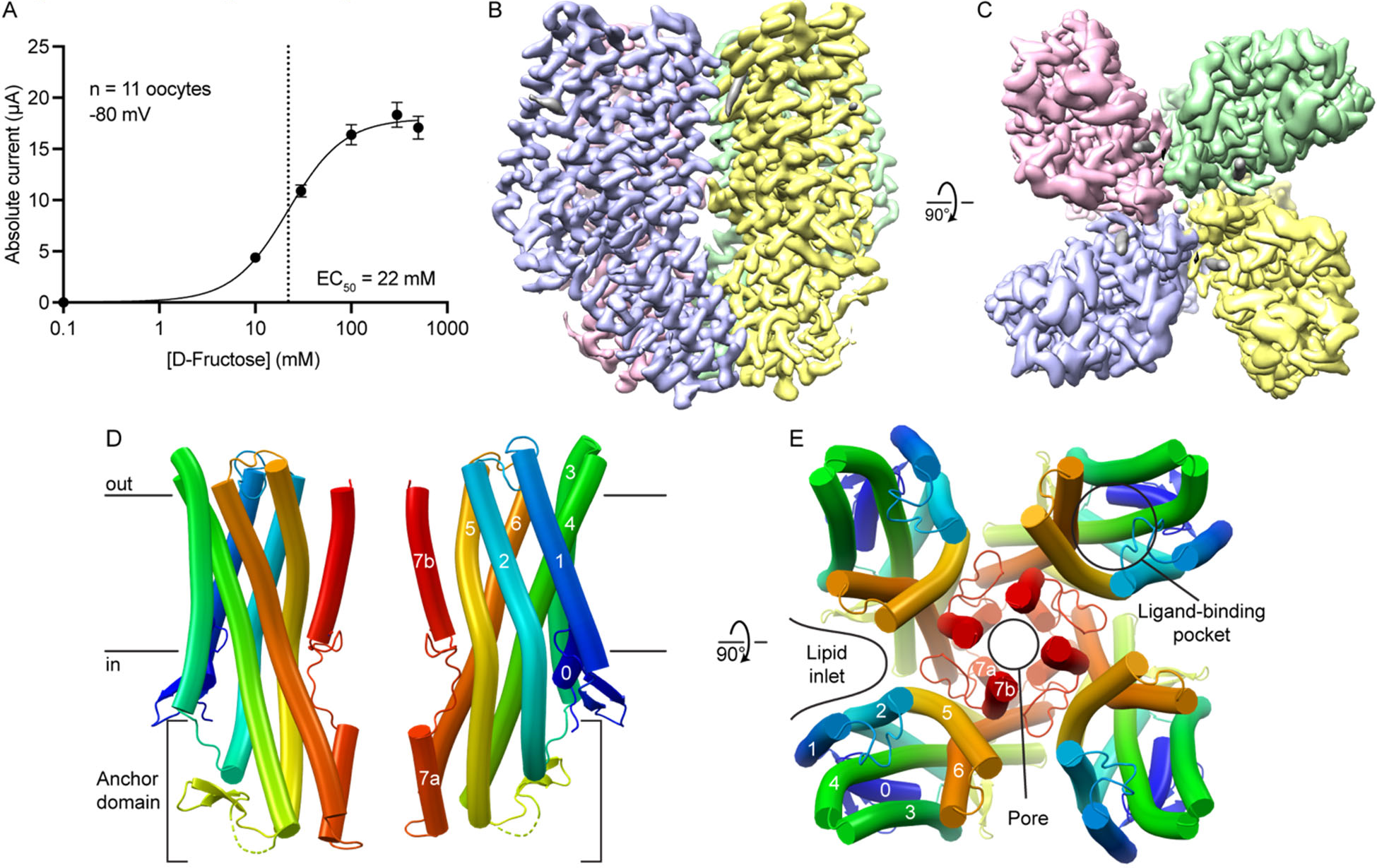
Cryo-EM structure of a tetrameric insect gustatory receptor, BmGR9. (A) D-Fructose dose-response profile of twin-strep-tagged BmGR9 expressed in *Xenopus* oocytes. The currents were measured at −80 mV (n = 11; data are shown as mean ± SEM), and the resulting EC_50_ value is 22 mM. (B-C) Surface representation of the Coulomb potential map of agonist-free BmGr9 (2.85 Å resolution; contour level of 0.24, around the protein structure), viewed from the membrane plane (B) and extracellular side (C). Each subunit of the tetramer is colored differently. (D) Cartoon representation of two opposing subunits from the BmGr9 tetramer viewed from the membrane plane. Black horizontal lines indicate the membrane boundaries. The intracellular anchor domain is indicated and the helices of one subunit are labeled. (E) Cartoon representation of the BmGr9 tetramer viewed from the extracellular side. The location of the pore and the fructose-binding pocket are indicated and the helices of one subunit are labeled. See also Figures S1-S5.

As anticipated from its homology to ORs, each BmGr9 subunit contains an N-terminal intracellular S0 helix and seven transmembrane helices (S1 to S7) with S7 broken into S7a and S7b (Figure 1D). BmGr9 is a tetramer with C4 symmetry, with the ion pore on the central four-fold axis (Figure 1E). The BmGr9 structure encompasses most of the protein, except for a disordered intracellular loop between S4 and S5 (residues 232-269), and the N and C termini (15 and 5 residues, respectively). Helices S1 and S3 only span the thickness of the lipid bilayer, while S2, S4, S5, S6, and S7a extend into the intracellular space to form an anchor domain similar to that observed in AbOrco and MhOr5 (Figure 1D; ^24,25^). Helix S7b contributes to the pore and helices S0-S6 expand outward from the central pore axis to form the ligand-binding pocket (Figure 1E). The main intersubunit contacts are around the transmembrane pore and the intracellular anchor domain (Figure 1D-E), burying a total of ∼2200 Å^2^ of surface area per subunit with ∼965 Å^2^ from the pore region and ∼1235 Å^2^ from the anchor. The four anchor domains thus form a tight intracellular bundle whereas deep lipid inlets largely isolate each subunit in the membrane plane (Figure 1E).

### BmGr9 and MhOr5 have distinct structural features

Both AbOrco and MhOr5 assemble as tetramers with a central ion pore formed by the C-terminal transmembrane segment, S7b. The core of each individual subunit of MhOr5 has a ligand-binding site in the middle of the membrane bilayer plane that appears inaccessible to aqueous solvent, consistent with its selectivity for hydrophobic ligands. Based on the pairwise superpositions of individual subunits, the structure of BmGr9 is more distant from AbOrco and MhOr5 than they are from each other (root mean squared distance (RMSD) values of 4.7 Å, 5.1 Å, and 3.8 Å for the BmGr9-AbOrco, BmGr9-MhOr5, and AbOrco-MhOr5 pairs, respectively). The secondary structure elements of BmGr9 are well defined in our cryo-EM density maps (Figure S6A), and several of these elements differ in length and orientation relative to the membrane plane when compared to MhOr5 (Figure 2A-B).

**Figure 2.**
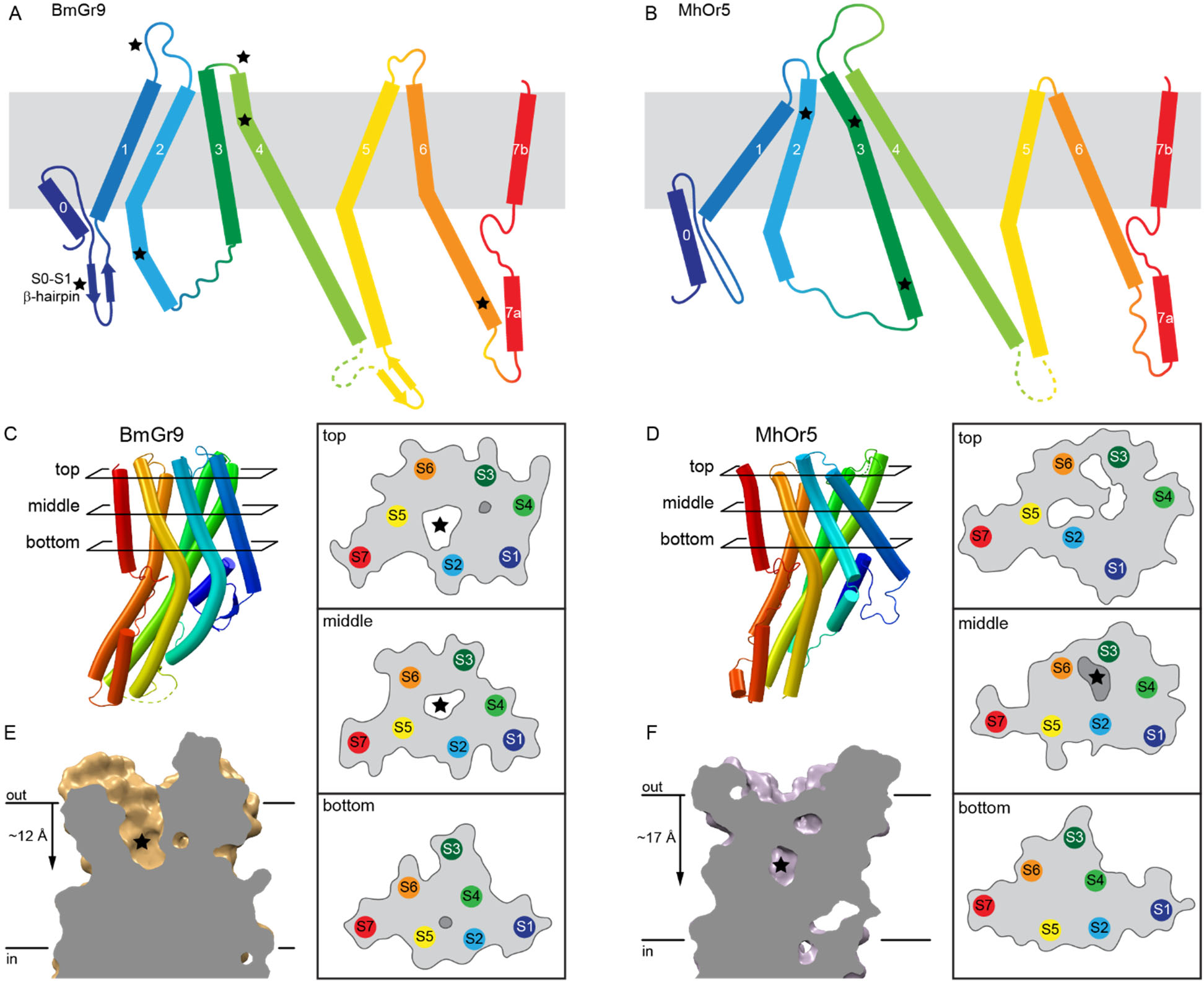
BmGr9 and MhOr5 have distinct structural features. (A-B) Diagrams comparing the topology of BmGr9 (A) and MhOr5 (B). Several of the features distinct between BmGr9 and MhOr5—such as loop and helix lengths, and kinks in helices—are highlighted with a star. (C-D) Transmembrane helix packing arrangements of BmGr9 (C) and MhOr5 (D) at different cross-sections (top, middle, bottom) across the membrane plane, as indicated on the cartoon representation of one subunit on the left-hand side. For each cross-section, the position of each transmembrane helix segment is illustrated, along with the protein surface boundary as a grey line. Internal cavities are marked in white when accessible to the extracellular aqueous solvent, and in darker grey for solvent-occluded cavities. Stars mark the fructose-binding pocket of BmGr9 (C) and the eugenol-binding pocket of MhOr5 (D). Of note, the S5 helix forms part of the fructose-binding pocket walls in BmGr9, but it is not in direct contact with the eugenol-binding pocket in MhOr5. (E-F) Vertical slices through the ligand-binding pocket in the transmembrane domain of one subunit of BmGr9 (E) and MhOr5 (F). The approximate position of the ligand is marked by a star, and the approximate distance from the extracellular membrane boundary to the bottom of the pocket is indicated on the left. See also Figure S6.

The most prominent structural differences between BmGr9 and MhOr5 are starred in Figure 2A-B. They include: (i) BmGr9 has a β-hairpin between the S0 and S1 helices, which neatly tucks under S0 (Figure 2C); (ii) The S1-S2 loop is longer in BmGr9 than in MhOr5 (10 versus 4 residues), and the S3-S4 loop—which closes access to the eugenol-binding pocket in MhOr5—is much shorter in BmGr9 than in MhOr5 (1 versus 28 residues); (iii) The BmGr9 S2 helix extends further into the intracellular space (7 residues longer than in MhOr5) and is more curved, with two kinks that lead the S2 helix to pack against the rest of the anchor domain; (iv) The S6 helix is longer in BmGr9 (45 versus 34 residues), with a sharp kink at the intracellular membrane boundary; and (v) the extracellular tip of the BmGr9 S4 helix has another sharp kink.

The combination of these five structural differences generates two global differences between BmGr9 and MhOr5. First, the differences in orientation and position of the S1-S6 helices within the plane of membrane (Figure 2C-D) and the lengths of the extracellular loops (Figure 2A-B) generate very different ligand-binding pockets (Figure 2E-F). BmGr9 has a ∼12-Å deep solvent-accessible ligand-binding pocket that totals ∼600 Å^3^ in volume, whereas MhOr5 has a much smaller pocket (∼100 Å^3^) that is located deeper in the membrane plane and is occluded from the extracellular solvent by its longer S3-S4 loop. Furthermore, S5 forms part of the fructose-binding pocket walls in BmGr9, unlike the S5 in MhOr5 which is not in direct contact with the eugenol-binding pocket (top and middle cross-sections in Figure 2C-D). Second, the presence of the S0-S1 β-hairpin and the longer S2 helix in BmG9 provide more extensive structural connections between the ligand-binding pocket formed by helices S1-S6 in the transmembrane domain and the intracellular anchor domain.

To assess whether structural features discussed in this study are linked to specific sequence signatures, we created separate family-wide sequence alignments of 1854 GRs and 3885 ORs (excluding ORCOs) from insects using a structure-based approach leveraging the AlphaFold Protein Structure Database^26^ (see methods for details). The sequence coverage of the two alignments is excellent, particularly at alignment positions corresponding to secondary structure elements (Figure S6B). From the aligned GR sequences, we extracted a subalignment of 74 DmGr43a subfamily members containing BmGr9 and the other experimentally validated fructose receptor orthologs, and we also used a previously published alignment of 176 ORCO sequences^24^.

Sequence covariation among the 1854 aligned GR sequences supports the idea that several of the structural features distinct to BmGr9 in comparison to MhOr5 are conserved in GRs, with 28 strongly evolutionarily coupled residue pairs (>90% confidence; Figure S6C-D). There is one evolutionarily coupled residue pair crosslinking the two strands of the N-terminal S0-S1 β-hairpin, suggesting that this β-hairpin is also conserved in GRs. A single pair is in the transmembrane region, corresponding to the buried salt bridge between E200 and R361 just below the base of the ligand-binding pocket. Intriguingly, this salt bridge is conserved in MhOr5 (D220 and R387), but not in AbOrco, suggesting that it is best conserved in subunits that naturally bind ligands. The twenty-six remaining pairs are all intrasubunit contacts within the anchor domain, and several of those link the extended intracellular portion of the S2 helix to S4 and S5. This suggests that the extended S2 helix (when compared to MhOr5; Figure 2A-B), and its packing against S4 and S5 to expand the anchor domain, is a conserved feature in GRs. This connection of the S2 helix to the anchor domain could provide additional leverage for allosteric communications between the ligand-binding pocket and other regions of the tetrameric ion channels, including the ion pore and its gate.

### Lipid headgroups penetrate the ion pore through intersubunit fenestrations

The cryo-EM map of agonist-free BmGr9 also contains strong density features that do not correspond to protein, especially in the lipid inlets between subunits (Figure 3A). We assigned several of these features to ordered phospholipids rather than detergent based on their well resolved bidentate shapes. Because no lipids were added during protein purification, we infer that these lipids originated from the HEK293 cell membrane and remained bound to the protein through sample preparation. We assigned these densities as phosphatidylcholine (PC; the most common lipid in HEK293 cells^27^), although the headgroups are unmodeled due to missing density.

**Figure 3.**
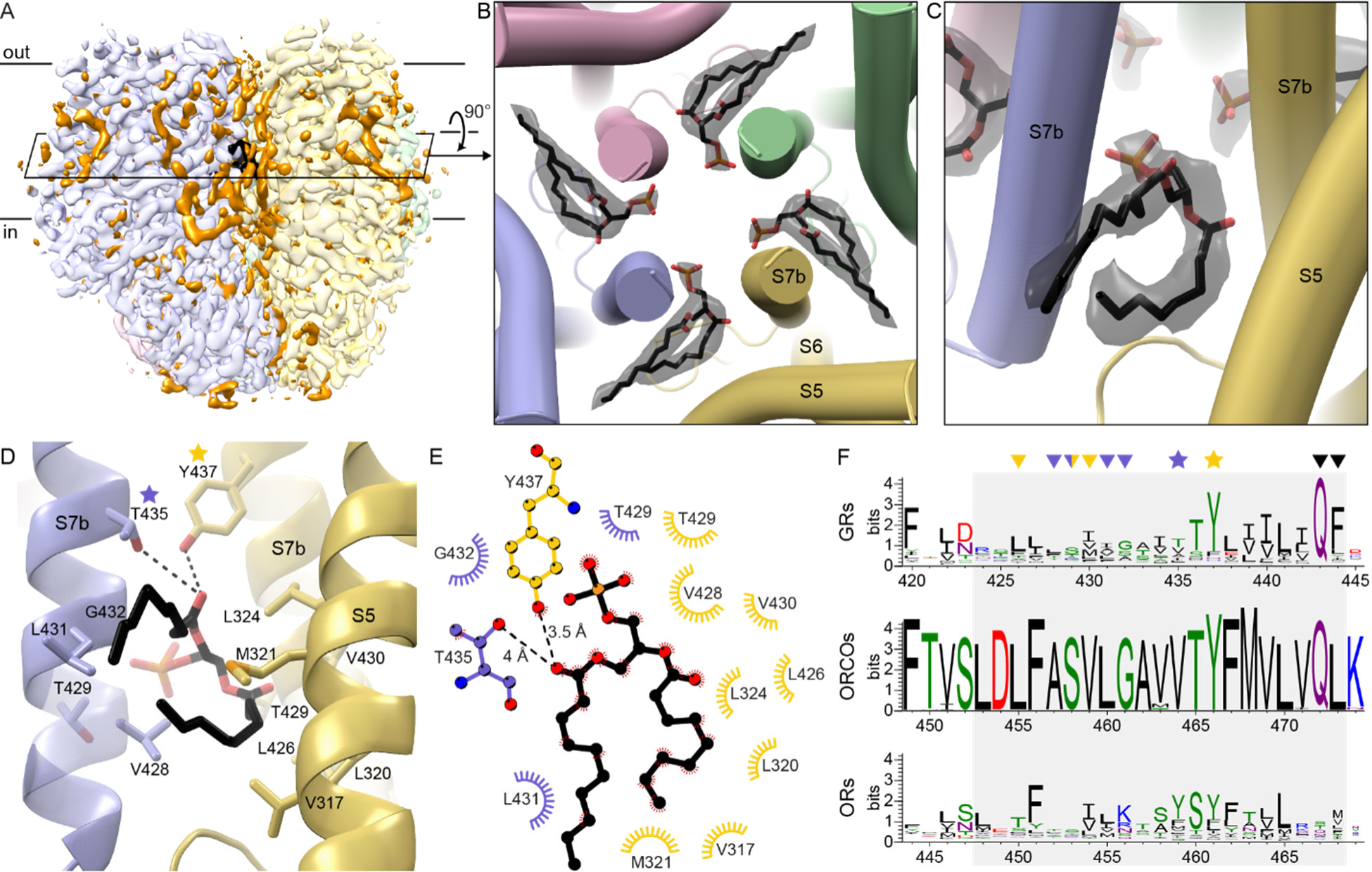
Lipid headgroups penetrate the BmGr9 pore through intersubunit fenestrations. (A) Cryo-EM map of agonist-free BmGr9 including non-protein densities (contour level 0.254). Each protein subunit of the tetramer is colored in a different pastel color, densities corresponding to ordered lipid or detergent molecules are orange, except for those of the pore-penetrating lipids which are colored black. The cross-section illustrated in (B) is indicated by the black outline. (B) Transmembrane cross-section of BmGr9 (viewed from extracellular side) highlighting densities (black transparent surface) for lipid with headgroups penetrating into the pore. Modeled lipids are shown as sticks (carbon in black, oxygen in red, phosphorus in orange). Only the phosphate of the lipid headgroup is modeled as the rest is disordered and lacks density. (C) View from the membrane plane of pore-penetrating lipid and corresponding cryo-EM density, highlighting its bidentate shape. (D) View of pore-penetrating lipid from membrane plane, with sidechains within 4.2 Å of the lipid shown as sticks. The lipid headgroup contacts the S7b helices from two BmGr9 subunits (blue and yellow, respectively), whereas the lipid interactions with the yellow S5 helix are hydrophobic contacts with a lipid tail. One turn of the blue S7b helix is transparent so that T429 is visible. Two highly conserved polar residues are starred. (E) Schematic of the interactions between a pore-penetrating lipid and two BmGr9 subunits (blue and yellow, respectively). Atoms are colored as in (D). The diagram was made with LigPlot+^41^. (F) Sequence logos of the helix S7b positions (grey box) of alignments of 1854 insect GR sequences, 176 ORCO sequences, and 3885 insect OR sequences, colored as follows: green, polar amino acids (G,S,T,Y,C,Q,N); blue, basic (K,R,H); red, acidic (D,E); and black, hydrophobic (A,V,L,I,P,W,F,M)). Residues interacting with the pore-penetrating lipids are marked with stars (polar residues) or arrowheads (aliphatic residues) colored according to panels (D) and (E). The pore-gating residues are marked by black arrowheads. See also Figure S7.

Four ordered lipids (one per subunit) project their head group from within the hydrophobic region of the bilayer into the aqueous ion pore, filling fenestrations in the pore walls at subunit interfaces (Figure 3B-C). The lipid enters the pore between S7b helices of adjacent subunits at the level of the inner leaflet. Interestingly, while associated lipids were not reported for AbOrco and MhOr5, analogous fenestrations capable of accommodating a lipid are observed in both structures (Figure S7A-B). Furthermore, bidentate densities consistent with such a pore-penetrating lipid are also observed in the map of our fructose-bound BmGr9 structure, as well as in the maps reported for eugenol-bound and DEET-bound MhOr5 (Figure S7C-E). The position of these pore-penetrating lipids suggests that they could contribute to the structural stability of the ion pore upon tetramerization, influence ion conduction and selectivity, or both.

In BmGr9, T435 and Y437 form polar contacts with the headgroup of the pore-penetrating lipid (Figure 3D-E). For Y437, tyrosine is highly conserved at this position across GRs (80.7%; Figure 3F) and ORCOs (99.4%; position 466 in AbOrco), and corresponds to the second residue of the GR family signature motif T**Y**hhhhhQF^28^ (where h is any hydrophobic amino acid). Tyrosine is also common at the corresponding position in ORs (52.9%; position 461 in MhOr5). For T435, small polar residues are common at this position in GRs (34.6% S, T, or N), while a tyrosine is common in ORs (51.1%), preserving the potential to hydrogen-bond with the lipid headgroup. The hydrophobic tails of the pore-penetrating lipids interact with non-polar sidechains from S7b and S5. As expected from their membrane-embedded positions, hydrophobic residues are enriched at each of these positions in GRs, ORCOs, and ORs (Figure 3D-F).

In summary, three observations suggest that the presence of pore-penetrating lipids is a conserved feature across both the GR and OR families: all available structures have fenestrations between S7b helices of adjacent subunits; lipid-shaped densities are present in similar positions in maps of both BmGr9 and MhOr5 structures, and the general properties of the lipid-interacting residues are conserved at the corresponding positions in sequence alignments of GRs and ORs.

### Pore gate opening involves an interaction network conserved between GRs and ORs

The BmGr9 pore has the same quadrivial architecture as AbOrco^24^ and MhOr5^25^ with a singular extracellular path down the center of the tetramer that opens into a large vestibule in the center of the membrane plane, then diverges into four lateral conduits formed between subunits (Figures 4A and S8A-D). The density for the central pore helix S7b and its sidechains is excellent in both the agonist-free and fructose-bound BmGr9 maps (Figure S6). In the agonist-free structure, starting from the extracellular face of the channel, the pore begins with a closed double-layer hydrophobic gate formed by F444 and I440 at the C terminus of S7b (Figure 4A-C). Like L473 in AbOrco and V468 in MhOr5 (Figure S8E), the F444 sidechains from each BmGr9 subunit protrude into the channel lumen to close the ion pore down to a diameter of 2.5 Å, creating a hydrophobic plug (Figure 4A-C). Below the hydrophobic gate, two threonines (T429 and T436) form two rings of hydroxyls that can stabilize cations (Figure 4C). AbOrco and MhOr5 also have hydroxyls in the same positions (Figure S8E). T436 is highly conserved across insect GRs—it is the first residue of the **T**YhhhhhQF signature motif. T429 is less conserved although predominantly polar or charged (71%, 100%, 74% across GRs, ORCOs and ORs respectively; Figure S8F).

**Figure 4.**
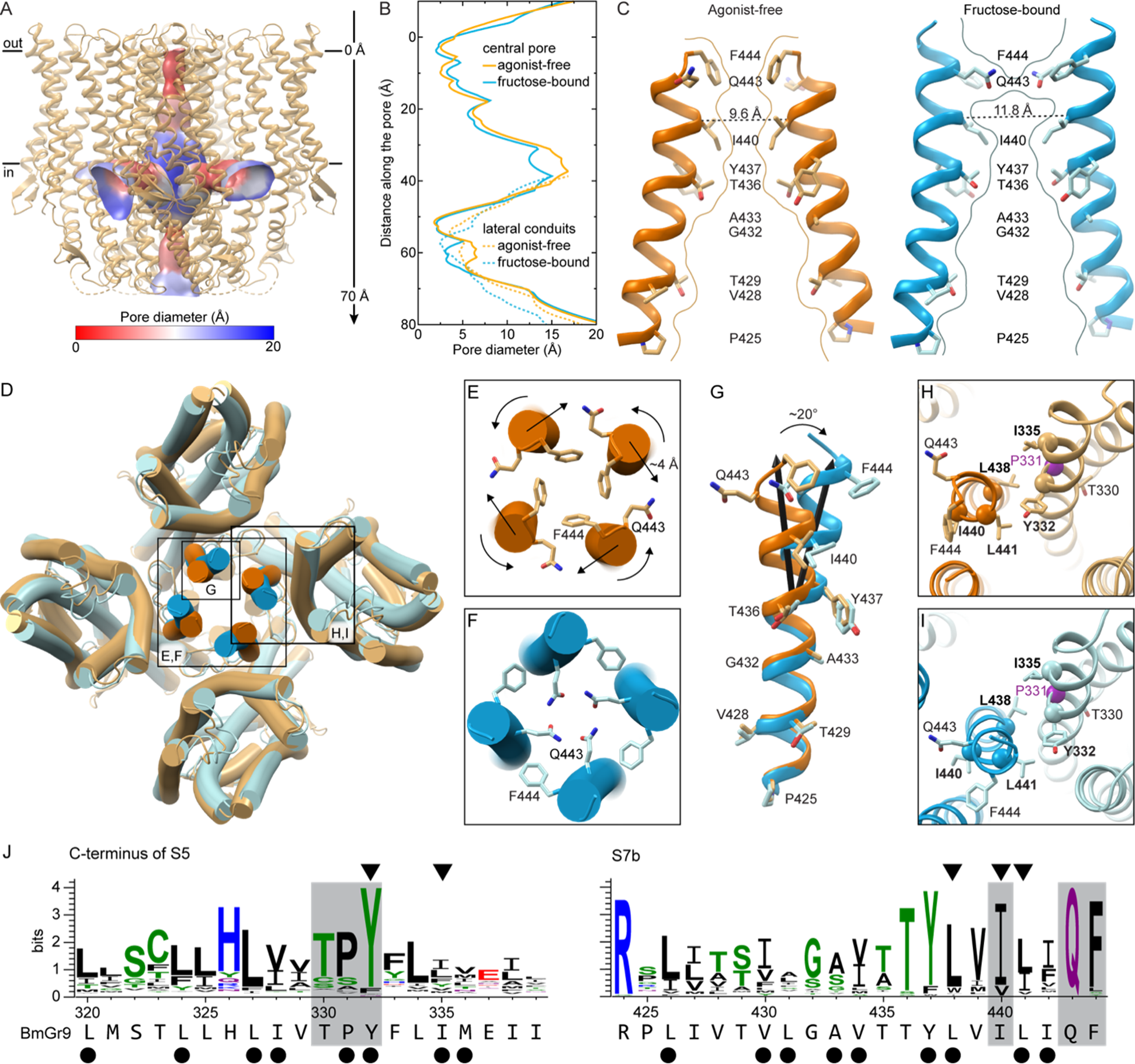
Fructose binding induces pore opening in BmGr9 through concerted motions of a conserved network of residues. (A) Cartoon representation of agonist-free BmGr9 with its quadrivial pore illustrated as a surface colored according to its diameter. Black lines make the membrane boundaries. The vertical axis for panel (B) is indicated on the right-hand side. (B) Diameter of the central ion conduction pathway (solid lines) and the lateral conduits (dashed lines) of agonist-free (orange) or fructose-bound (blue) BmGr9. The y axis shows the distance from the outer membrane boundary toward the intracellular space for the central pore, and the distance along the conduit for the lateral conduits. (C) Cartoon representation of two opposing S7b helices with the sidechains of pore-lining residues in sticks for agonist-free (left) and fructose-bound (right) BmGr9. The pore surface at the central vertical cross-section is marked by curved lines. F444 creates a hydrophobic seal in agonist-free BmGr9, whereas Q443 creates a hydrophilic ring more suitable for cation conduction in fructose-bound BmGr9 although the pore diameter is largely unchanged. The Cα-to-Cα distances between the two opposing hydrophobic gate residues I440 are indicated and marked by dashed black lines. (D) Superposition of agonist-free (orange) and fructose-bound (blue) BmGr9. The inset panels E-I are indicated. (E-F) Zoomed-in view of from the extracellular side of the agonist-free (E) and fructose-bound (F) BmGr9 central pore with conserved gating residues Q443 and F444 shown as sticks. The twisting motion of the gating residues and displacement of the C termini of the S7b helices upon fructose binding are indicated by arrows. (G) One S7b helix viewed from the center of the ion pore, from the superimposed structures of the agonist-free (orange) and fructose-bound (blue) BmGr9 tetramer. The pore-lining sidechains are shown as sticks and labeled. Black arrows mark the central axes of the top half of the helices, highlighting the 20° kink and displacement upon fructose binding. (H-I) Zoomed-in view of the interaction between helices S5 and S7b in the agonist-free (H) and fructose-bound (I) BmGr9 structures. P331 is marked by a purple Cα sphere. Mutations to alanine of the residues labeled in bold and marked by the other Cα spheres increase basal currents^30^. Pore-gating Q443 and F444, and the conserved T330 residue facing the fructose-binding pocket are also shown as sticks for reference. (J) Sequence logos of the C-terminal half of S5 and of S7b for the alignment of Gr43a subfamily members (74 sequences). The BmGr9 sequence is indicated below. Residues forming the hydrophobic S5-S7b interface are marked by black dots, and locations of the alanine mutations that increase basal currents are marked by arrowheads. The conserved TPY motif in S5 and pore-gating residues in S7b are highlighted in grey. See also Figure S8.

Located immediately below F444 near the extracellular opening of the ion pore, Q443 points away from the pore in the agonist-free structure but rotates into the ion pathway in the fructose-bound structure (Figure 4C-F). The Q443 sidechains thus create a hydrophilic ring more suitable for cation conduction in the fructose-bound structure, although the pore diameter is largely unchanged (Figure 4B). In addition to its twisting along the helix axis, the C terminus of S7b also moves out from the center of the tetramer by ∼4 Å. T436, near the pore-penetrating lipid headgroup, acts as a fulcrum for these movements, such that only the portion of S7b above T436 moves (Figure 4G). As a result, the pore widens the most at the other hydrophobic constriction, with the Cα-to-Cα distance between opposing I440 residues increasing from 9.6 Å to 11.8 Å, and the pore diameter from 2.5 Å to 3.2 Å (Figure 4B-C).

Q443 and F444 are highly conserved in GRs and ORCOs; they are the last two residues of the GR family signature TYhhhhh**QF** motif^28^ (Figure S8F). In BmGr9, the signature motif adheres to the GR consensus: 436-TYLVILIQF-444. This motif is notably less conserved in ORs, most of which form heteromers with ORCO^8,20^; however, this glutamine is conserved in homotetrameric MhOr5, which is from a basal insect species that lacks an ORCO ortholog^29^. This Q467 in MhOr5 also swings into the pore upon agonist-induced channel opening and mutating it to alanine or arginine eliminated agonist-activated calcium influx, suggesting that this rearrangement is critical for receptor function^25^.

Mutations of the hydrophobic gate residues in BmGr9 (I440 and F444) also support a role for both residues in gating^30^. I440 mutations (I440A and I440Q) increased the inward resting current and decreased the responsiveness to fructose compared to wildtype, suggesting that the hydrophobic gate has been disrupted, allowing cations to pass through the pore without agonist binding^30^. The F444A mutation increased the EC_50_ for fructose and reduced the fructose-induced currents, suggesting a change in open-closed equilibrium or conductance^30^.

Pore helix S7b is tightly packed with—and its movements coupled to—helix S5, which connects directly to the binding pocket in BmGr9 (Figure 4H-J). BmGr9 residues L441 on S7b and Y332 on S5 correspond to MhOr5 residues L465 and Y362, respectively, two highly conserved and evolutionarily coupled hydrophobic amino acids in ORs that also move in concert between the agonist-free and eugenol bound MhOr5 structures^25,31^. The Y362A mutation in MhOr5 impaired eugenol activation while Y362F showed no effect, supporting the importance of the aromatic group at this position^31^. BmGr9 residue Y332 is highly conserved in GRs while L441 is conserved as an aliphatic residue (Figure 3F). Mutations to alanine of either Y332, L441, or two other conserved hydrophobic residues at the S7b-S5 interface—L438 and I335—increased the resting current of BmGr9^30^, suggesting that the tight packing of S7b and S5 is required for proper ligand gating.

The S5-S7b interaction connects the pore to the binding pocket, suggesting that small fructose-induced movements in S5 could gate the pore. Y332 is part of a highly conserved TPY motif in the Gr43a subfamily (Figure 4J). T330 faces the ligand-binding pocket and likely interacts with the fructose (see also below). P331 breaks the backbone hydrogen-bonding pattern on S5, and thus could reduce the energy barrier to local rearrangements and thus facilitate allosteric communication between the ligand-binding pocket and the pore.

### Fructose-binding pocket is lined with polar and aromatic residues

In contrast to the solvent-inaccessible eugenol-binding cavity observed in MhOr5^25^, BmGr9 has a deep solvent-accessible cavity at the analogous location, with its opening on the extracellular face near the center of each subunit (Figure 2, S9A-B). In each subunit, the extracellular termini of helices S0-S6 splay out, with helices S2-S5 forming the walls of this deep pocket. At the bottom of this pocket, the fructose-bound map contains additional density when compared to the agonist-free map (Figure 5A-B). While the resolution of the map at this location is not high enough to resolve the fructose binding pose, the location is consistent with published mutational analysis^30^. Furthermore, computational docking suggests that a fructose molecule is readily accommodated in this pocket. That is, when docking a β-D-fructopyranose (the most abundant form of D-fructose in solution^32^), the most energetically favored positions are just adjacent to the observed density (Figure S9C). Upon real-space refinement of the protein model and docked fructose against the cryo-EM map, the fructose roughly fits inside the density (Figure S9C). We are therefore confident in assigning the bottom of the deep pocket as the fructose-binding site, and discuss it as such below, although we refrain from describing specific fructose-protein interactions.

**Figure 5.**
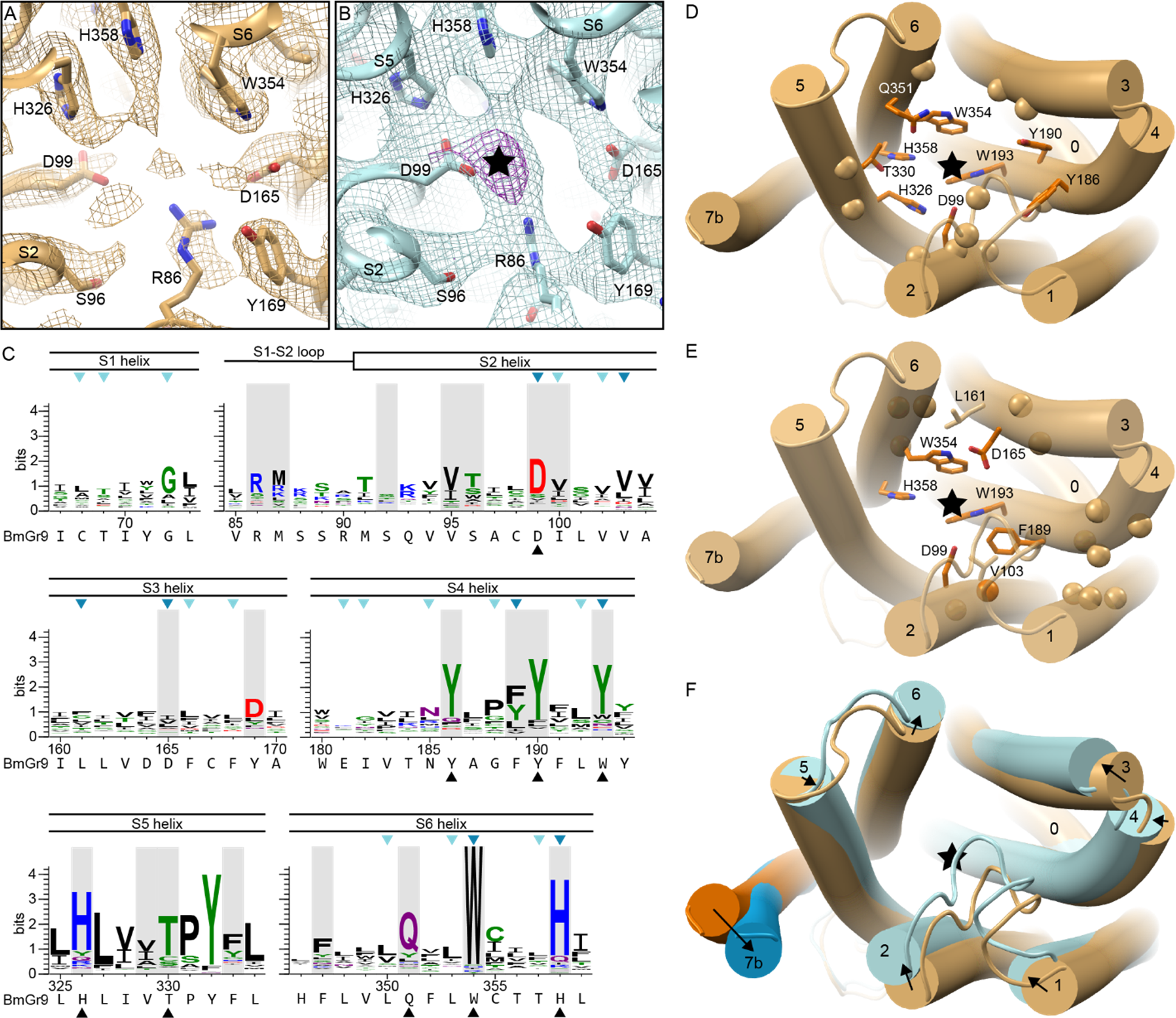
BmGr9 ligand-binding pocket and its fructose-induced conformational changes. (A) Cryo-EM density around the ligand-binding site for the agonist-free BmGr9 map contoured at 3.5σ. Nearby sidechains are shown as sticks and labeled. (B) Cryo-EM density around the ligand-binding site for the agonist-free BmGr9 map contoured at 4.5σ (cyan). The black star marks strong density not observed in the agonist-free map shown in panel A, as indicated by the presence of a 10σ density feature (magenta). (C) Sequence logos from the Gr43a subfamily alignment covering motifs participating in the ligand-binding pocket, with the corresponding secondary structure elements indicated above, and the BmGr9 sequence included below each logo. The 21 residues with surface accessible to the pocket are shaded grey, and black arrowheads mark highly conserved pocket-facing residues highlighted in panel D. Blue arrowheads mark positions assessed by alanine substitutions in BmGr9^30^; light blue showed no change in fructose response, bright blue showed reduced response to fructose. (D-F) Cartoon representations of one BmGr9 subunit viewed from the extracellular side, with the black star marking the approximate fructose position. (D) Residues participating in the binding pocket are shown as sidechain sticks for highly conserved residues or marked by Cα spheres for the others (corresponding to black arrowheads and grey highlights in panel C, respectively). (E) Positions previously assessed by alanine substitutions in BmGr9^30^. Mutations to the eight sidechains shown in sticks eliminated fructose-induced currents, while mutations at the 15 other positions marked by spheres had no effect on fructose-induced currents. Positions facing the ligand-binding pocket are dark orange, others are in light orange. The cartoon representation transparency reveals otherwise hidden spheres. (F) Superimposed individual subunits of agonist-free (orange) and fructose-bound (blue) BmGr9 with black arrows highlighting the movement of helices from agonist-free to fructose-bound. See also Figure S9.

Nine of the 21 residues on the fructose-binding pocket surface are highly conserved in the Gr43a subfamily (>66% identity; Figure 5C-D). As described above, the TPY motif on S5 connects the binding pocket to the pore movements (Figures 4H and 5D). S5 residue T330, along with H326, faces the binding pocket. Both are highly conserved and are near the other highly conserved polar and aromatic residues in the bottom half of the ligand-binding pocket (S2 residue D99; S4 residues Y186, Y190, and W193 (68.6% Y and 80% aromatic); and S6 residues Q351, W354, and H358). Of the 21 residues we include as forming the pocket, 7 have been previously investigated by mutagenesis^30^. Of those, conserved residues D99, W193, W354, and H358, as well as D165 and F189 (70% conserved as an aromatic residue), are all essential for BmGr9 function, as mutating each residue individually to alanine dramatically reduced or eliminated fructose-activated channel function^30^ (Figure 5E). Two additional residues (of 16 tested) adjacent to but not on the surface of the binding pocket, V103 and L161 (81.7% and 58.0% aliphatic, respectively), were also important for the fructose response (Figure 5E); they may structurally enable the allosteric communication between the ligand-binding site and the pore gate. The overall high level of sequence conservation in the pocket (9 of the 45 most conserved residues in the BmGr9 subfamily), particularly at the bottom of it, suggests that maintaining aromatic and hydrophilic sidechains is important for ligand binding and is an important first step to convert fructose binding to pore opening.

In both a superposition of the agonist-free and fructose-bound BmGr9 subunit structures (Figure 5F) and an alignment-free distance difference matrix (Figure S9D), we observe relative movements of helices S2-S6 that reshape the binding pocket upon agonist binding (Figures 5F and S9A-B). S2—with its highly conserved D99 residue closest to the cryo-EM density assigned to fructose—moves the most, and most helices move inward towards the central axis of the protein (Figure 5F). This coordinated movement of the helices could convey changes caused by fructose binding to the gates of the ion pore.

### Ligand-binding pocket comparison across the GR family

Using the experimentally determined BmGr9 structure as a reference point, we explored the potential of bioinformatic approaches to inform structure-function relationships among GRs more broadly across the GR family. To assess evolutionary relationships among GRs, we created a phylogenetic tree from our structure-based multiple sequence alignment of 1854 GRs and 41 ORs (Figures 6A and S10). As chemical detection is a primary function of these receptors, we focused our analysis on the putative ligand-binding pockets of members of four subfamilies that contain members with relatively well-established chemosensory functions: the Gr5a and Gr43a subfamilies, which mediate sugar sensing; the Gr63a subfamily, which mediates CO_2_ sensing; and the ORs, which mediate responses to volatile organic compounds (Figures 6A and S11).

**Figure 6.**
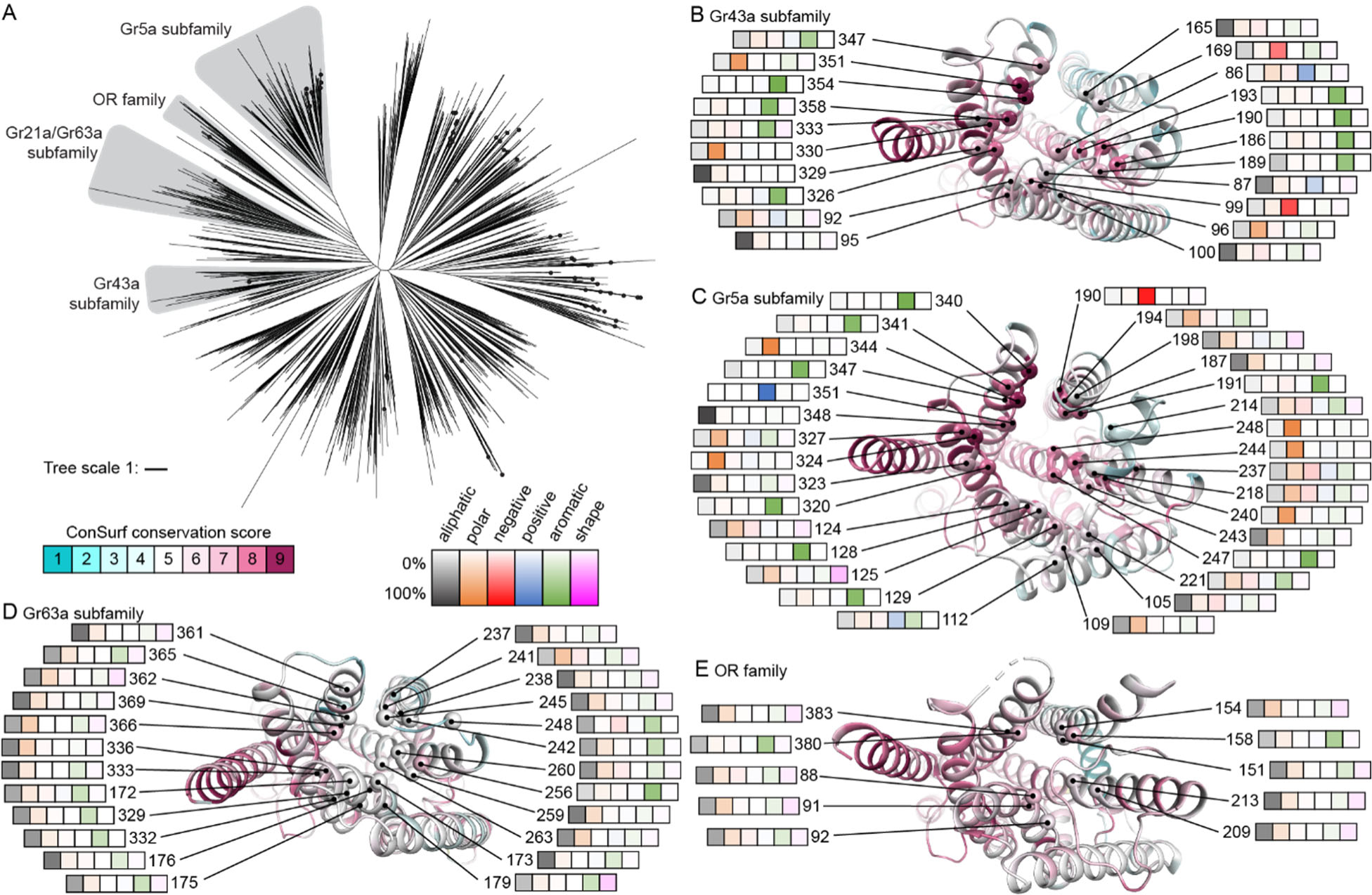
ligand pocket comparison across GR family. (A) Phylogenetic tree of 1895 insect GR sequences with subfamilies of interest highlighted in grey. The tree includes a clade of 41 OR sequences. Drosophila GRs are represented by black dots. (See also Figures S10-11.) (B-E) Cartoon representations of single subunits of a representative subfamily member viewed from the extracellular side. The secondary structure cartoons are colored by ConSurf^29^ score based on the sequence alignment of the corresponding GR subfamily. The Cα carbons of residues forming part of the predicted ligand-binding pocket and crevice are represented as a sphere. Each of these residues is labeled with its residue number in the reference protein and a heatmap capturing the frequency of different amino acid types at that position in the aligned sequences of the corresponding GR subfamily. In each heatmap, from left to right are aliphatic amino acids (Ala, Cys, Leu, Met, Val; black), polar (Asn, Gln, Ser, Thr; orange), negatively charged (Asp, Glu; red), positively charged (Arg, Lys; blue), aromatic (His, Phe, Trp, Tyr; green), or shape-determining (Gly, Pro; pink) residues. The following subunits and corresponding protein (sub)families are illustrated: (B) Gr43a subfamily (74 sequences) on the agonist-free BmGr9 structure; (C) Gr5a subfamily (251 sequences) on the AlphaFold2 model of Gr5a (UniProt ID: Q9W497); (D) Gr63a subfamily (107 sequences) on the AlphaFold2 model of Gr63a (UniProt ID: Q9VZL7); and (E) the OR family (3885 sequences) on the agonist-free MhOr5 structure (PDB ID: 7LIC). (See also Figures S10-S13.)

Experimentally determined structures are available for an OR (MhOR5) and for BmGr9 (a Gr43a ortholog), but not for Gr5a or Gr63a subfamily members, although AlphaFold predictions are available. We assessed the ability of AlphaFold2 to provide accurate structural predictions of GR and OR family members by comparing the monomeric models of BmGr9 and AbOrco from the AlphaFold database^26^ to their experimentally determined agonist-free structures (the database has no MhOr5 model). While neither structure was in the AlphaFold2 training set, both models correspond well to the respective structures (RMSD values of 0.9 Å for BmGr9 and 0.3 Å for AbOrco). We thus used the AlphaFold models of *D. melanogaster* Gr5a and Gr63a to identify the residues likely to form the surfaces of their putative ligand-binding pockets (Figure S12). Using ConSurf^29^, we mapped sequence conservation across 251 Gr5a and 107 Gr63a subfamily members to the corresponding representative AlphaFold model, and of 74 Gr43a subfamily members and 3885 ORs to the structures of BmGr9 and MhOr5, respectively (Figure 6B-E, S12).

While the Gr5a and Gr43a subfamilies are evolutionarily divergent (Figure 6A), both contain receptors involved in sugar detection, raising the possibility of similarities in their ligand-binding pockets. Indeed, despite their distinct evolutionary histories, their putative ligand-binding pockets are similarly enriched for aromatic and polar amino acids. Among Gr43a subfamily members, nine of the 21 pocket-lining positions are most commonly occupied by aromatic amino acids, along with four by polar, two by acidic, one by basic, and five by aliphatic amino acids (Figure 6B). Among Gr5a subfamily members, 11 of the 31 pocket-lining residues are most commonly occupied by aromatic amino acids, 12 by polar, one by acidic, one by basic, and six by aliphatic amino acids (Figure 6C). Of the most highly conserved positions (ConSurf score ≥8), six are aromatic, two polar and one negatively charged in the Gr43a subfamily (Figure 6B), while seven are aromatic, seven polar, two aliphatic, one positively and one negatively charged in the Gr5a subfamily (Figure 6C). This abundance of aromatic, polar, and charged amino acids is consistent with the role of aromatic and hydrophilic residues in sugar binding in other proteins^33,34^. As detailed below, it is not shared by Gr63a subfamily members or ORs.

Despite similarities in the chemical character of the binding pockets between the Gr43a and Gr5a subfamilies, their pocket sizes differ dramatically (Figure S12A-B, E). The predicted Gr5a binding pocket is much larger than the pocket we observe in the BmGr9 structure (∼2100 Å^3^ versus ∼600 Å^3^; Figure S12E). The larger pocket in Gr5a arises from several key predicted structural differences: Gr5a’s longer S3-S4 loop forms a V-shape with two short helices that enable S3 and S4 to splay further from the subunit center to enlarge the pocket, while its shorter S1-S2 loop further widens the exposed opening (Figure S13B). These structural features of Gr5a are present across the subfamily, with shorter S1-S2 and longer S3-S4 loops a distinct feature of Gr5a subfamily members compared to other GRs or to ORs (Figure S13A; p<0.01 Steel-Dwass test). These larger pockets could accommodate a wider variety of sugars, consistent with the ability of gustatory neurons expressing Gr5a subfamily members to respond to polysaccharides as well as monosaccharides^11^ (Figure S12).

The Gr63a subfamily forms another divergent clade of GRs, whose members can form CO_2_ sensors^1,13^ (Figure 6A). In contrast to the sugar receptors above, aliphatic amino acids predominate in the putative ligand-binding pockets of Gr63a subfamily members, with 20 of 24 pocket-lining positions mostly occupied by aliphatic amino acids, and just three by aromatic, and one by polar amino acids (Figure 6D). Similarly, for ORs, which are primarily involved in the detection of volatile organic compounds, 8 of the 10 pocket-lining positions are most commonly occupied by aliphatic amino acids, with the remaining two positions mostly occupied by aromatic amino acids (Figure 6E). The pocket sizes of representatives of both of these subfamilies are substantially smaller than those of the sugar receptors (∼150 Å^3^ for the Gr63a AlphaFold model and ∼100 Å^3^ for the MhOr5 structure; Figure S12C-E). Thus, receptors implicated in detecting different classes of chemicals differ in both pocket size and chemical composition, consistent with key roles in determining ligand specificity.

## DISCUSSION

Our work provides an initial view of the structural features that underlie the chemoreceptor activity of a member of the insect gustatory receptor family. The structure of the BmGr9 homotetramer bears substantial resemblance to the distantly related olfactory receptor MhOr5, including its quadrivial channel architecture. However, BmGr9 contains an additional S0-S1 β-hairpin and a longer S2 helix that together provide more extensive structural connections between the anchor domain and the transmembrane regions than in MhOr5. BmGr9’s ligand-binding pocket also differs from that in MhOr5, as the BmGr9 pocket is much larger and involves additional receptor surfaces, is lined with aromatic and polar residues rather than hydrophobic residues, and is open to the extracellular milieu. These distinct characteristics provide a structural basis for the distinct functions of the two receptors, with MhOr5 acting as an olfactory receptor for volatile, hydrophobic compounds like eugenol, and BmGr9 acting as a gustatory receptor for fructose, a water-soluble carbohydrate. Sequence analyses indicate that these distinctive features are conserved across other ORs and Gr43a subfamily members. At a mechanistic level, fructose binding to BmGr9 elicits small movements of the S1-S6 helices that surround the ligand-binding pocket. These movements narrow the pocket, with the movement of S5, which interacts with both the bound fructose and the S7b pore helix, potentially promoting the pore-opening motion of S7b.

Although GRs exhibit substantial sequence diversity, they do share a characteristic sequence motif near their C-termini: TYhhhhhQF^8^. In the BmGr9 structures, we observe a clear function associated with each of the 4 most highly conserved amino acids present in this motif on the pore-forming helix S7b. The TY pair shapes the structure and hydrophilic character of the ion pathway: the four T436 sidechains form a hydrophilic ring below the second hydrophobic gate—formed by the middle hydrophobic residue (I440) in the conserved motif—while the Y437 sidechains interact with pore-penetrating phospholipid headgroups. The QF pair plays a role in gating. The F444 sidechains from each of the four subunits form a hydrophobic plug at the extracellular pore opening in the closed agonist-free state. These phenylalanines swing out of the way upon fructose binding, as the adjacent Q443 sidechains swing into the pore, making the opening hydrophilic. The conservation of the sequences in this motif across the GR family suggests that helix S7b’s contributions to BmGr9 function will be conserved in other GRs.

### Pore-penetrating lipids as a common feature of GRs and ORs

Based on the available experimental structural data for BmGr9, AbOrco^24^, and MhOr5^25^, as well as the conservation of the TY sequence motif in GRs, ORs and ORCOs, the pore-penetrating lipids we observe bound to BmGr9 are likely a feature conserved across the superfamily. Fenestrations in the pore walls are observed in all available structures and density consistent with a lipid is present in both agonist-free and fructose-bound BmGr9 structures, suggesting that the lipids form part of a stable pore structure. Fenestrations opening the ion path to the membrane environment have been observed in several other ion channel families, including ionotropic glutamate receptors, cys-loop receptors, voltage-gated potassium channels and sodium channels, and mechanosensitive two-pore domain potassium (K2P) channels^35–39^. In some channels, such fenestrations are transient, while in others they are observed in both closed and open channel states. In many cases these fenestrations allow activators or blockers to enter the pore—including some endogenous ligands, and they have been implicated in the mechanisms of well-established drugs including common local and general anesthetics^39,40^. The fenestrations in insect GR and OR channels could therefore also represent opportunities for chemical modulation of their activity.

### Ligand-binding pockets of evolutionarily divergent sugar-sensing GRs share common features

The BmGr9 ligand-binding pocket in our structures differs significantly from a previously reported homology model, which was not based on AlphaFold^30^. Only ∼7 of the 21 residues that participate in the pocket in our experimentally determined structure are shared with the earlier prediction^30^. Observed contributions of residues from the S5 helix and the S1-S2 extracellular loop to the pocket were not predicted in the earlier model, and the pocket-lining residues in the S2, S3, S4 and S6 helices largely differ. Strikingly, six of the eight mutations from that study that had strong effects on fructose-induced currents^30^ face the experimentally determined binding pocket. In contrast, of the 11 mutations with no effect^30^, none face the experimentally determined binding pocket. These mutagenesis results suggest that residues distributed around the pocket are important for fructose binding and channel gating. This likely enables the exquisite selectivity of BmGr9 for fructose among other monosaccharides^17^.

Our bioinformatics analyses leveraging AlphaFold, which more accurately models BmGr9, indicate that the ligand-binding pockets of Gr5a subfamily members—a second set of GRs involved in sugar sensing—share characteristics with those of the Gr43a subfamily. As the two subfamilies are otherwise evolutionary divergent, this likely reflects convergence between these families of sugar receptors. That the pocket is larger in the Gr5a subfamily also suggests a structural explanation for the responsiveness of gustatory neurons expressing Gr5a subfamily members to di- and poly- as well as monosaccharides^11^, whereas BmGr9 and its orthologs like Gr43a are fructose-specific. Finally, our bioinformatics analyses suggest the members of the Gr63a subfamily of CO_2_ receptors have highly hydrophobic binding pockets (Figure 6). This suggests that CO_2_ detection could involve hydrophobic interactions with pocket residues or binding of a hydrophobic ligand or co-factor. Taken together, this work illustrates how the combination of experimental structures and structure-based sequence analyses can help delineate the evolution and function of large protein families like the insect GRs and ORs.

Expanding these comparisons to different GRs subfamilies with well-defined ligand classes, the predicted ligand-binding pocket dimensions and chemical properties are shaped by structural features that are conserved within subfamilies but differ between subfamilies with distinct ligand types. For example, G5a subfamily members have large and polar predicted pockets strongly suggesting that they are indeed di- or polysaccharide-gated ion channels, consistent with their role in sugar detection in sensory neurons. Our work thus illustrates that the combination of experimental structures and structure-based sequence analyses can help delineate the evolution and function of large protein families like the insect GRs and ORs.

### Limitations of this study

At the structural level, although agonist binding to BmGr9 promotes structural changes conducive to ion flow, the ion path appears too narrow for ion conduction in the structures captured here. In the agonist-free state, the extracellular entrance to BmGr9’s ion pore is sealed by a double-layered hydrophobic gate formed by helix S7b. Upon agonist binding, S7b movements replace the outer layer of the gate’s hydrophobic plug (formed by an F444 from each subunit) with hydrophilic residues (Q443), and they also increase the distance between the hydrophobic residues (I440) of the gate’s inner layer, dilating the pore by ∼1.8 Å. However, agonist binding does not dramatically alter the narrowest constriction of the ion pore, with an oxygen-to-oxygen distance of ∼5.4 Å at the level of Q443 in fructose-bound BmGr9. This contrasts with the ∼9.2 Å oxygen-to-oxygen distance reported for the analogous Q467 position in agonist-bound MhOr5^25^. This suggests that additional pore conformations exist for BmGr9. Structure determination in a lipid environment, such as in nanodiscs, rather than in detergent, could reveal these additional conformational shifts. Finally, it will also be of interest to examine the positioning of the pore-penetrating lipids in a lipid environment.

While our bioinformatic analyses explore GRs beyond BmGr9, the depth of analysis is constrained by the paucity of functional information about most GRs. Mechanistic studies of GRs have proven challenging, with most attempts to study GRs in heterologous expression systems failing to observe channel activity (BmGr9 is an exception). While genetic evidence suggests that many GRs participate in heteromeric complexes, which GRs hetero-oligomerize and which GRs participate in ligand binding or serve primarily structural roles remain to be established. A second limitation of our bioinformatic analyses is that while AlphaFold2 predictions agree well with the experimentally determined structures of AbOrco and BmGr9, this does not guarantee that all predictions will be as accurate. However, the presence of shared, and potentially convergent, features in the ligand-binding pockets of evolutionarily divergent sugar receptors, and their contrast with receptors for other classes of chemicals, supports the utility of this approach for capturing structural features on a scale not readily achieved by experimental determination alone.

## Acknowledgements

We thank Elizabeth J. May for help with the development of the HEK293-based protein expression system, José A. Velilla, Shamayeeta Ray, Gerardo E. Zavala and Samuel P. Berry for sharing technical expertise, and Leslie Griffith and members of the Garrity and Gaudet labs for discussions and comments on the manuscript. Funding was provided by National Institute of Deafness and Other Communication Disorders grant R21DC018497 (P.A.G. and R.G.), National Institute of General Medical Sciences grants R01GM120996 (R.G.) and R01GM096053 (D.L.T), National

Institute of Allergy and Infectious Diseases R01AI157194 (P.A.G.), NSF Simons center at Harvard Simons postdoctoral award (S.W.), Charles A. King Trust Postdoctoral Research Fellowship, Bank of America, N.A., Co-Trustees (W.J.L.), and a Warren Alpert Foundation Distinguished Scholar Award (W.J.L.). We acknowledge the use of resources from the Harvard Cryo-EM Center for Structural Biology.

## Author contributions

H.M.F., R.M.W., W.J.L., P.A.G. and R.G. designed experiments. H.M.F. performed molecular biology, protein expression and purification, cryo-EM sample preparation, cryo-EM data collection, model building and refinement, and structural analyses under the supervision of R.M.W. and R.G. R.M.W. supervised the collection of and analyzed the cryo-EM data. W.J.L. performed molecular biology and oocyte electrophysiology. H.M.F., S.W., D.L.T., P.A.G. and R.G. designed and performed bioinformatic and statistical analyses. H.M.F., S.W., R.M.W., W.J.L., P.A.G. and R.G. created the figures. H.M.F., S.W., P.A.G. and R.G. wrote the paper, with input from all authors.

## STAR Methods

### RESOURCE AVAILABILITY

#### Lead contact

Further information and requests for resources and reagents should be directed to and will be fulfilled by the lead contact, Rachelle Gaudet.

#### Materials availability

Plasmids generated are available from the lead contact with a completed materials transfer agreement.

#### Data and code availability

- The cryo-EM maps have been deposited to the Electron Microscopy Data Bank (EMDB) (accession numbers: EMD-43129 and EMD-43130, respectively, for the agonist-free and fructose-bound maps), and the refined coordinates to the Protein Data Bank (PDB IDs: 8VC1 and 8VC2, respectively, for the agonist-free and fructose-bound BmGr9 structures). The GR family sequence alignment and phylogenetic tree have been deposited XXXX. All other data is available from the corresponding authors upon reasonable request.
- This paper does not report any original code.
- Any additional information required to reanalyze the data reported in this paper is available from the lead contact upon request.

### EXPERIMENTAL MODEL AND SUBJECT DETAILS

The *E. coli* DH5α strain, cultured in LB medium (Thermo Fisher Scientific) at 37°C, was used to amplify plasmids. Human embryonic kidney (HEK) 293F inducible GnTI-suspension cells (Thermo Fisher Scientific, A39242) were cultured in Expi293 expression medium (Thermo Fisher Scientific) at 37°C, supplied with 8% CO_2_. HEK293T cells were cultured in Dulbecco’s modified Eagle’s medium (DMEM, Corning) supplemented with 10% (v/v) fetal bovine serum (Corning), 1X GlutaMAX (Gibco), and 100 U/mL penicillin-streptomycin (Lonza) at 37°C and 5% CO_2_. *Xenopus laevis* oocytes (EcoCyte Bioscience) were cultured at 18°C in ND96 media containing (mM): 2 KCl, 96 NaCl, 2.0 MgCl_2_, 1.8 CaCl_2_, 5 HEPES-NaOH pH 7.4 supplemented with penicillin/streptomycin.

### METHOD DETAILS

#### Constructs

For protein expression and cryo-EM, the BmGr9 sequence (gift from Kazushige Touhara)^17^ was inserted into the pHR-CMV-TetO2_3C-Twin-Strep lentiviral expression plasmid with a IRES EmGFP reporter (Addgene ID: 113883) that was modified to introduce an N-terminal Twin-Strep tag (WSHPQFEKGGGSGGGSGGSAWSHPQFEK). For electrophysiology experiments, the same N-terminally tagged BmGr9 open reading frame was inserted into pOX (Addgene ID: 3780) using Gibson assembly.

#### Stable polyclonal cell line for BmGr9 expression

BmGr9 was expressed using a lentivirus expression system to create a stable polyclonal cell line in the Expi293F inducible GnTI-cell line (ThermoFisher Scientific) using published protocols^42^. Briefly, lentiviral particles were produced by co-transfecting 1.8×10^7^ HEK293T cells seeded 24 hours prior with 16 µg of the lentiviral expression plasmid described above, 16 µg psPAX2 packaging plasmid (Addgene ID: 12260), and 16 µg pMD2.G envelope plasmid (Addgene ID: 12259) using polyethylenimine (Polysciences). The lentivirus-containing supernatant was harvested after 3 days, then applied to 10×10^6^ Expi293F inducible GnTI-cells and incubated for 3 days to allow for genomic integration and establishment of a polyclonal stable cell line. Transduced cells were expanded and frozen in aliquots for long-term storage and use.

#### Protein expression and purification

The stably transduced polyclonal cell line was grown at 37°C with 8% carbon dioxide in Expi293 expression medium (ThermoFisher Scientific) to 3×10^6^ cells/mL, then induced with 5 μg/mL doxycycline and 5 mM sodium butyrate for 72 hours. Cells were collected by centrifugation at 600 g for 10 min, washed with phosphate buffered saline (PBS; Corning), resuspended in lysis buffer (20 mM Tris pH 8.25, 150 mM NaCl, 1 μM pepstatin A, 1 mM phenylmethylsulfonylfluride (PMSF), 1 mM benzamidine), and lysed using an Avestin Emulsiflex C5 with 4 passes at 5-15 kpsi. The lysate was cleared using a 15 min low-speed spin (9,700 g) and membranes were pelleted at 185,000 g for 2 hours, flash frozen, and stored at −80°C. All purification steps were performed at 4°C. Thawed membranes from 2 L of culture were homogenized in 120 mL solubilization buffer (PBS, 1 μM pepstatin A, 1 mM PMSF, 1 mM benzamidine, and 2% (w/v) n-dodecyl β-D-maltoside (DDM; Anatrace)) using a glass Potter-Elvehjem grinder, then rocked for 1.5 hr at 4°C. Detergent-insoluble material was pelleted at 185,000 g for 40 min and the supernatant was loaded onto a 1-mL Strep-Tactin XT 4Flow cartridge at 1 mL/min at 4°C. The resin was washed 5 x 2 column volumes (CV) of wash buffer (100 mM Tris pH 8.0, 150 mM NaCl, 1 mM EDTA, 0.05% (w/v) DDM, 1 mM DTT) then 6 x 0.5 CV of elution buffer (wash buffer with 50 mM biotin). To elute BmGr9, the column was rocked for 30 min with 3 CV of elution buffer and collected by displacing the column with 1 additional CV. The rocking elution was repeated 4-5 times until all protein was eluted. Elutions containing BmGr9 were combined and further purified using size exclusion chromatography (SEC) on a Superose 6 10/300 GL (Cytiva) equilibrated with SEC buffer (20 mM Tris-HCl pH 8.25, 150 mM NaCl, 1 mM DTT, 1 mM EDTA, 0.01% glyco-diosgenin (GDN; Anatrace)). For the fructose-bound structure the SEC buffer also contained 270 mM D-fructose. BmGr9-rich fractions were combined, concentrated to 3.2-3.3 mg/mL with a 100 kDa molecular weight cut-off centrifugal filter (Millipore), and used immediately for preparing cryo-EM grids.

#### Cryo-EM sample preparation

Concentrated BmGr9 protein (3 μL) was deposited onto 400 mesh Quantifoil Cu 1.2/1.3 grids that had been glow discharged in a PELCO easiGLOW (Ted Pella) at 0.39 mBar, 15 mA for 30 s. Agonist-free samples were vitrified in 100% liquid ethane using a Vitrobot Mark IV (Thermo Fisher Scientific), with a wait time of 0 s, blot time of 3 s, drain time of 0 s, and a blot force of 0 at 100% humidity. Fructose-bound samples were prepared with a blot time of 9 s.

#### Cryo-EM data collection and processing

Cryo-EM data were collected on a 300 kV Titan Krios G3i Microscope (Thermo Fisher Scientific) equipped with a K3 direct electron detector (Gatan) and a GIF quantum energy filter (20 eV) (Gatan) using counted mode at the Harvard Cryo-Electron Microscopy Center for Structural Biology at Harvard Medical School. Data were acquired utilizing image shift and real-time coma correction by beamtilt using the automated data collection software SerialEM^43^; nine holes were visited per stage position acquiring two movies per hole. Details of the data collection and dataset parameters are summarized in Table 1. Dose-fractionated images were gain normalized, aligned, dose-weighted and summed using MotionCor2^44^. Contrast transfer function (CTF) and defocus value estimation were performed using CTFFIND4^45^. Details of the data processing strategy are shown in Figure S2. In short, particle picking was carried out using crYOLO^46^) followed by 3D classification within Relion^47^. For the fructose-bound sample, a second round of 3D classification was performed to further improve the particle set (Figure S4). The selected particles were then subjected to Bayesian polishing following 3D refinement with C4 symmetry imposed. The data then underwent CTF refinement and nonuniform refinement with C4 symmetry imposed, in cryosSPARC^48^. Outputs from nonuniform refinement underwent CryoSPARC Local Refinement to produce the final 2.85 Å (3.23 Å C1) and 3.98 Å (6.4 Å C1) reconstructions for the Apo and Fructose samples respectively. Structural biology applications used in this project were compiled and configured by SBGrid^49^.

**Table 1:**
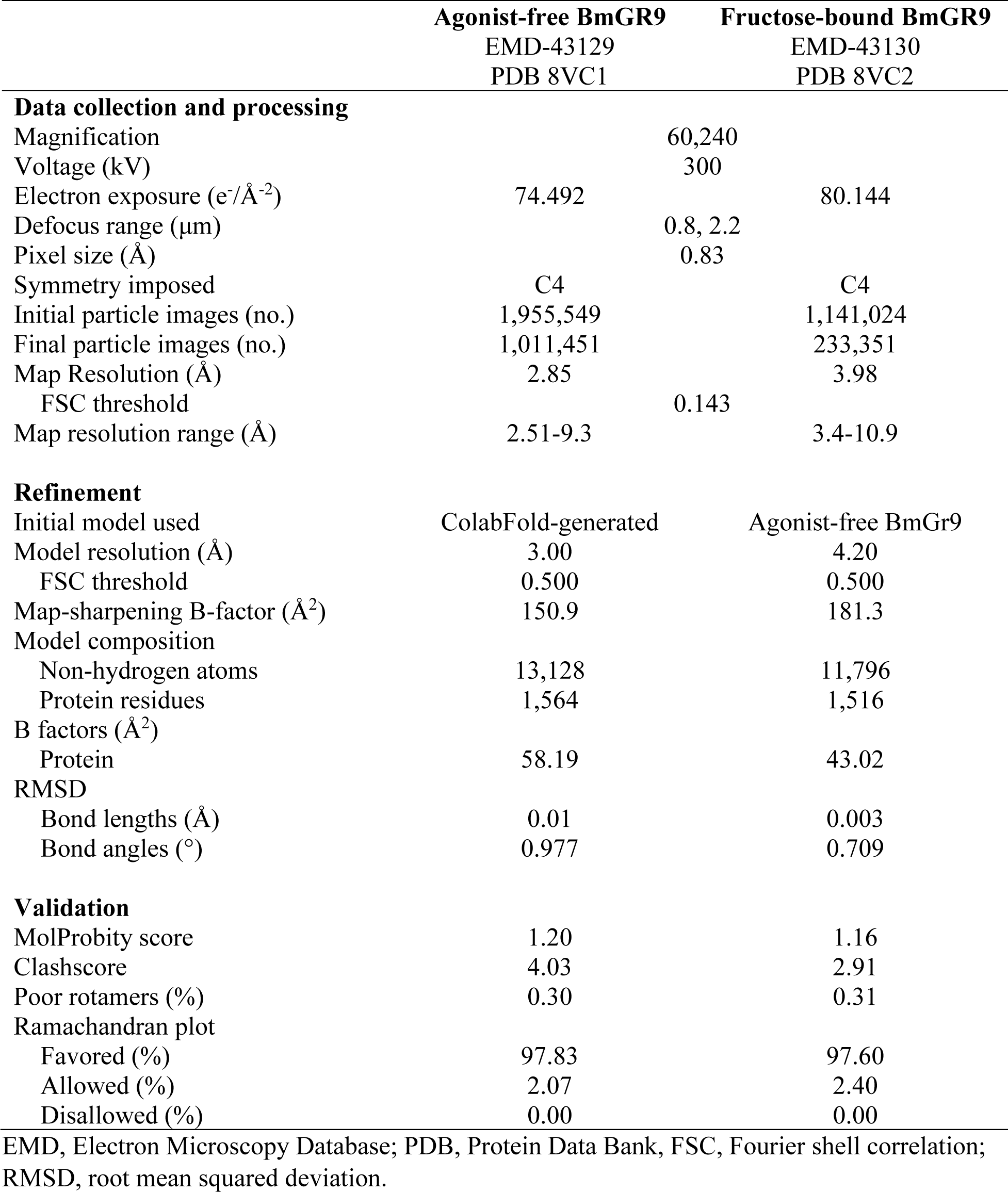
Cryo-EM data collection, refinement, and validation statistics.

#### Model building and refinement

A monomeric model of BmG9 was generated using ColabFold^50^. The rank 1 model was placed in the map with 4-fold symmetry using DockInMap in PHENIX^51^, and refined through cycles of manual rebuilding in Coot^52^, real-space refinement in PHENIX with macrocycles including morphing, global minimization, nhq_flips, and ADP, under secondary structure and NCS constraints, and remodeling by simulations run in the ISOLDE plugin of ChimeraX^53^. The refinement statistics are summarized in Table 1.

#### Electrophysiology

*Xenopus laevis* oocytes (EcoCyte Bioscience) were injected with 20 ng cRNA and cultured at 18°C in ND96 media containing (mM): 2 KCl, 96 NaCl, 2.0 MgCl_2_, 1.8 CaCl_2_, 5 HEPES-NaOH pH 7.4 supplemented with penicillin/streptomycin. Electrophysiological recordings were performed two days after injection by two-electrode voltage clamp using a OC-725 amplifier. Whole-cell currents were elicited by 2.5-s voltage ramp from −150 to +90 mV from a holding potential of −80 mV in ND96 (pH 7.4), filtered at 1 kHz, and recorded in pCLAMP 8 software (Molecular Devices).

#### Generation of multiple sequence alignments

The members of the GR family of proteins display a high degree of functional and sequence diversity which makes their sequence alignment and functional classification a difficult task. To explore the sequence-function relationship of GRs, we first created a seed alignment of 57 different GR sequences from *D. melanogaster* by structurally aligning monomer models downloaded from the AlphaFold Protein Structure Database^26^ using MUSTANG^54^. Using this seed alignment, we created a profile hidden Markov model (HMM) using the hmmbuild tool from the HMMER software package (hmmer.org) and searched the Uniref50 database (release-2023_02) for sequences matching this profile HMM using the hmmsearch tool in HMMER. We filtered the set of sequences obtained in the previous step to select sequences with bit score larger than 50, sequence length between 300 and 600, and from species belonging to the class insecta. Next, we added the sequences of known fructose-activated GRs to this filtered set of sequences. We also added sequences of the two odorant receptor proteins with published structures, AbOrco and MhOr5, to generate the final set of 1895 sequences. We then aligned these sequences to the profile HMM created in the first step.

We created a multiple sequence alignment of insect ORs using a method similar to the one we used for creating the alignment of GRs. We created a profile HMM from structural alignment of 62 Alphafold2 models of *D. melanogaster* ORs and identified insect sequences of 300-600 amino acids in length. Some of these sequences turned out to be ORCO proteins which we removed from this set and added the sequence for MhOr5. Finally, we aligned this set of sequences to the profile HMM that was created in the first step to create an alignment of 3885 OR sequences.

We used the previously published alignment of 176 ORCO sequences^24^ without any modifications.

#### Phylogenetic tree construction and classification

Using IQ-TREE 2^55^, we inferred 20 separate maximum likelihood phylogenetic trees from the alignment of 1895 GR sequences with the Le-Gascuel (LG) substitution model^56^, optimized equilibrium frequencies, and across-site rate variation using the discrete gamma model with 8 categories. We selected the tree with the best log likelihood as the final tree for further analysis. The resulting tree with aBayes posterior probability branch support values is in Figure S10, and the annotated tree in Figure S11. Based on the position of the AbOrco and MhOr5 sequences and annotations of nearby sequences, we determined that one small clade in the tree contained ORs (41 sequences), marking the branch point of divergence of ORs from the GR family. These 41 sequences were removed from the ‘all GR’ set of 1854 sequences for further analyses.

For the next part of our analysis, we assumed that GRs that appear in the same subfamily of the tree to be functionally related to each other. This assumption was supported by the fact that all known D-fructose receptors—DmGr43a, BmGr9, AgGr25, *Helicoverpa armigera* Gr4, and *Apis mellifera* Gr3—clustered in the same subfamily. We extracted the corresponding set of 74 sequences, which we refer to as the Gr43a subfamily. To functionally identify other subfamilies in the tree, we looked for *D. melanogaster* GRs in each subfamily and assigned the function of the *D. melanogaster* GRs found in that subfamily to the entire subfamily. Using this strategy, we extracted two more subfamilies in addition to Gr43a subfamily for further analysis (Figure S11): The Gr63a subfamily containing known CO_2_ receptors (107 sequences) and the Gr5a subfamily containing many known sugar receptors (251 sequences).

#### Sequence conservation analysis

For the OR family and each of the three GR subfamilies chosen for further analysis, we selected a structural model of a representative GR: the agonist-free MhOr5 structure (PDB ID: 7LIC) for the OR family; the agonist-free BmGr9 structure for the Gr43a subfamily; and the AlphaFold2 models of Gr5a (UniProt ID: Q9W497) and Gr63a (UniProt ID: Q9VZL7) for the Gr5a and Gr63a subfamilies, respectively. For each of the five corresponding sequences alignments, we performed sequence conservation analysis using ConSurf^29,57,58^, using the WAG evolutionary model.

#### Ligand-binding pocket analysis

The predicted ligand-binding pockets for the Gr5a and Gr63a subfamilies were identified in the central region of the transmembrane S1-S6 bundle of each representative Alphafold2 model. We confirmed that the approximate size and location of each predicted pocket was a reasonable representative by comparing them to 4-5 additional AlphaFold2 models chosen from different branches of the same subfamily. The experimentally determined ligand-binding pocket of BmGr9 was used for the Gr43a subfamily and of MhOr5 for the OR family. All residue positions with solvent-exposed atoms facing the respective pockets were selected (Figure S12). The observed amino acid frequencies at each position were computed from the respective sequence alignment. The heatmaps in Figure 6 were obtained by summing the amino acid frequencies according to the following amino acid groupings based on chemical properties: aliphatic (Ala, Cys, Leu, Met, Val; black), polar (Asn, Gln, Ser, Thr; orange), negatively charged (Asp, Glu; red), positively charged (Arg, Lys; blue), aromatic (His, Phe, Trp, Tyr; green), and shape-determining (Gly, Pro; pink) residues.

#### Determination of loop lengths

Despite high diversity and low sequence similarity of GR proteins, our structure-based and profile HMM-assisted approach produced an alignment with high overlap (>80%) coverage for the transmembrane helix regions (Figure S6B). Assuming that sequence regions corresponding to transmembrane helices are well aligned with each other, the poorly aligned regions between two transmembrane helices must correspond to the loops that connect those helices. Because we have the experimentally determined structure of BmGr9, we used its sequence as a reference to define boundaries of helices S1 to S7 for all sequences in the GR alignment and the selected subfamily alignments (helix boundaries for S1 are residues 52-79; S2, 90-135; S3, 141-176; S4, 178-229; S5, 287-339; S6, 345-389; S7a, 395-409; S7b, 424-444). For the OR alignment, we similarly used the MhOr5 structure as a reference (S1, 49-76; S2, 78-123; S3, 130-159; S4, 198-251; S5, 317-368; S6, 371-415; S7a, 420-434; S7b, 448-472). To determine the loop lengths, we counted the number of residues connecting the two adjacent transmembrane helices in each sequence, after shortening each helix by 1 residue on each end to allow for noise in the alignment. To avoid including incomplete sequences, we only counted sequences that had non-zero loop length.

#### Pore dimension analysis

We used the HOLE program^59^ to analyze the dimensions of ion conduction pores of agonist-free and fructose-bound structures of BmGr9.

#### Fructose docking

We used AutoDock Vina^60,61^ to dock β-D-fructopyranose in the fructose-bound structure. The AutoDock search space was set to a cuboid of 25 x 30 x 40 Å^3^ spanning the length and width of a monomer. We then used real-space refinement in PHENIX^51^ (macrocycles including morphing, global minimization, nhq_flips, and ADP, under secondary structure and NCS constraints) with the AutoDock rank 1 position of fructose which modeled the fructose into the density.

### QUANTIFICATION AND STATISTICAL ANALYSIS

For the electrophysiology experiments described in Figure 1A and S1A-D, n refers to independent biological replicates. Shapiro-Wilk tests were used to assess the normality of all datasets (p≤0.05 rejected normal distribution). Nonparametric tests consisted of Kruskal-Wallis followed by a Steel-Dwass post hoc test for multiple comparisons (JMP11, SAS).

## References

1. Liman, E.R., Zhang, Y.V., and Montell, C. (2014). Peripheral coding of taste. Neuron 81, 984–1000. 10.1016/j.neuron.2014.02.022.

2. Stork, N.E., McBroom, J., Gely, C., and Hamilton, A.J. (2015). New approaches narrow global species estimates for beetles, insects, and terrestrial arthropods. Proc Natl Acad Sci U S A 112, 7519–7523. 10.1073/pnas.1502408112.

3. Rosenberg, Y., Bar-On, Y.M., Fromm, A., Ostikar, M., Shoshany, A., Giz, O., and Milo, R. (2023). The global biomass and number of terrestrial arthropods. Sci Adv 9, eabq4049. 10.1126/sciadv.abq4049.

4. Potts, S.G., Imperatriz-Fonseca, V., Ngo, H.T., Aizen, M.A., Biesmeijer, J.C., Breeze, T.D., Dicks, L.V., Garibaldi, L.A., Hill, R., Settele, J., and Vanbergen, A.J. (2016). Safeguarding pollinators and their values to human well-being. Nature 540, 220–229. 10.1038/nature20588.

5. World Health, O. (2014). A global brief on vector-borne diseases. World Health Organization. 2014. https://iris.who.int/handle/10665/111008.

6. Renault, D., Angulo, E., Cuthbert, R.N., Haubrock, P.J., Capinha, C., Bang, A., Kramer, A.M., and Courchamp, F. (2022). The magnitude, diversity, and distribution of the economic costs of invasive terrestrial invertebrates worldwide. Sci Total Environ 835, 155391. 10.1016/j.scitotenv.2022.155391.

7. Joseph, R.M., and Carlson, J.R. (2015). Drosophila Chemoreceptors: A Molecular Interface Between the Chemical World and the Brain. Trends Genet 31, 683–695. 10.1016/j.tig.2015.09.005.

8. Robertson, H.M. (2019). Molecular Evolution of the Major Arthropod Chemoreceptor Gene Families. Annu Rev Entomol 64, 227–242. 10.1146/annurev-ento-020117-043322.

9. Matthews, B.J., Dudchenko, O., Kingan, S.B., Koren, S., Antoshechkin, I., Crawford, J.E., Glassford, W.J., Herre, M., Redmond, S.N., Rose, N.H., et al. (2018). Improved reference genome of Aedes aegypti informs arbovirus vector control. Nature 563, 501–507. 10.1038/s41586-018-0692-z.

10. Wanner, K.W., and Robertson, H.M. (2008). The gustatory receptor family in the silkworm moth Bombyx mori is characterized by a large expansion of a single lineage of putative bitter receptors. Insect Mol Biol 17, 621–629. 10.1111/j.1365-2583.2008.00836.x.

11. Fujii, S., Yavuz, A., Slone, J., Jagge, C., Song, X., and Amrein, H. (2015). Drosophila sugar receptors in sweet taste perception, olfaction, and internal nutrient sensing. Curr Biol 25, 621–627. 10.1016/j.cub.2014.12.058.

12. Dweck, H.K.M., and Carlson, J.R. (2020). Molecular Logic and Evolution of Bitter Taste in Drosophila. Curr Biol 30, 17–30 e13. 10.1016/j.cub.2019.11.005.

13. McMeniman, C.J., Corfas, R.A., Matthews, B.J., Ritchie, S.A., and Vosshall, L.B. (2014). Multimodal integration of carbon dioxide and other sensory cues drives mosquito attraction to humans. Cell 156, 1060–1071. 10.1016/j.cell.2013.12.044.

14. Miyamoto, T., Slone, J., Song, X., and Amrein, H. (2012). A fructose receptor functions as a nutrient sensor in the Drosophila brain. Cell 151, 1113–1125. 10.1016/j.cell.2012.10.024.

15. Hoshino, R., Sano, H., Yoshinari, Y., Nishimura, T., and Niwa, R. (2023). Circulating fructose regulates a germline stem cell increase via gustatory receptor-mediated gut hormone secretion in mated Drosophila. Sci Adv 9, eadd5551. 10.1126/sciadv.add5551.

16. Kikuta, S., Endo, H., Tomita, N., Takada, T., Morita, C., Asaoka, K., and Sato, R. (2016). Characterization of a ligand-gated cation channel based on an inositol receptor in the silkworm, Bombyx mori. Insect Biochem Mol Biol 74, 12–20. 10.1016/j.ibmb.2016.04.010.

17. Sato, K., Tanaka, K., and Touhara, K. (2011). Sugar-regulated cation channel formed by an insect gustatory receptor. Proc Natl Acad Sci U S A 108, 11680–11685. 10.1073/pnas.1019622108.

18. Xu, P., Wen, X., and Leal, W.S. (2020). CO(2) per se activates carbon dioxide receptors. Insect Biochem Mol Biol 117, 103284. 10.1016/j.ibmb.2019.103284.

19. Benton, R., and Himmel, N.J. (2023). Structural screens identify candidate human homologs of insect chemoreceptors and cryptic Drosophila gustatory receptor-like proteins. Elife 12. 10.7554/eLife.85537.

20. Sato, K., Pellegrino, M., Nakagawa, T., Nakagawa, T., Vosshall, L.B., and Touhara, K. (2008). Insect olfactory receptors are heteromeric ligand-gated ion channels. Nature 452, 1002–1006. 10.1038/nature06850.

21. Wicher, D., Schafer, R., Bauernfeind, R., Stensmyr, M.C., Heller, R., Heinemann, S.H., and Hansson, B.S. (2008). Drosophila odorant receptors are both ligand-gated and cyclic-nucleotide-activated cation channels. Nature 452, 1007–1011. 10.1038/nature06861.

22. Yan, H., Jafari, S., Pask, G., Zhou, X., Reinberg, D., and Desplan, C. (2020). Evolution, developmental expression and function of odorant receptors in insects. J Exp Biol 223. 10.1242/jeb.208215.

23. Chen, Y.D., and Dahanukar, A. (2020). Recent advances in the genetic basis of taste detection in Drosophila. Cell Mol Life Sci 77, 1087–1101. 10.1007/s00018-019-03320-0.

24. Butterwick, J.A., Del Marmol, J., Kim, K.H., Kahlson, M.A., Rogow, J.A., Walz, T., and Ruta, V. (2018). Cryo-EM structure of the insect olfactory receptor Orco. Nature 560, 447–452. 10.1038/s41586-018-0420-8.

25. Del Marmol, J., Yedlin, M.A., and Ruta, V. (2021). The structural basis of odorant recognition in insect olfactory receptors. Nature 597, 126–131. 10.1038/s41586-021-03794-8.

26. Tunyasuvunakool, K., Adler, J., Wu, Z., Green, T., Zielinski, M., Zidek, A., Bridgland, A., Cowie, A., Meyer, C., Laydon, A., et al. (2021). Highly accurate protein structure prediction for the human proteome. Nature 596, 590–596. 10.1038/s41586-021-03828-1.

27. Dawaliby, R., Trubbia, C., Delporte, C., Noyon, C., Ruysschaert, J.M., Van Antwerpen, P., and Govaerts, C. (2016). Phosphatidylethanolamine Is a Key Regulator of Membrane Fluidity in Eukaryotic Cells. J Biol Chem 291, 3658–3667. 10.1074/jbc.M115.706523.

28. Robertson, H.M. (2015). The Insect Chemoreceptor Superfamily Is Ancient in Animals. Chem Senses 40, 609–614. 10.1093/chemse/bjv046.

29. Ashkenazy, H., Abadi, S., Martz, E., Chay, O., Mayrose, I., Pupko, T., and Ben-Tal, N. (2016). ConSurf 2016: an improved methodology to estimate and visualize evolutionary conservation in macromolecules. Nucleic Acids Res 44, W344–350. 10.1093/nar/gkw408.

30. Morinaga, S., Nagata, K., Ihara, S., Yumita, T., Niimura, Y., Sato, K., and Touhara, K. (2022). Structural model for ligand binding and channel opening of an insect gustatory receptor. J Biol Chem 298, 102573. 10.1016/j.jbc.2022.102573.

31. Hopf, T.A., Morinaga, S., Ihara, S., Touhara, K., Marks, D.S., and Benton, R. (2015). Amino acid coevolution reveals three-dimensional structure and functional domains of insect odorant receptors. Nat Commun 6, 6077. 10.1038/ncomms7077.

32. Alexandersson, E., and Nestor, G. (2022). Complete (1)H and (13)C NMR spectral assignment of d-glucofuranose. Carbohydr Res 511, 108477. 10.1016/j.carres.2021.108477.

33. Gabius, H.J., Andre, S., Jimenez-Barbero, J., Romero, A., and Solis, D. (2011). From lectin structure to functional glycomics: principles of the sugar code. Trends Biochem Sci 36, 298–313. 10.1016/j.tibs.2011.01.005.

34. Hudson, K.L., Bartlett, G.J., Diehl, R.C., Agirre, J., Gallagher, T., Kiessling, L.L., and Woolfson, D.N. (2015). Carbohydrate-Aromatic Interactions in Proteins. J Am Chem Soc 137, 15152–15160. 10.1021/jacs.5b08424.

35. Dong, Y.Y., Pike, A.C., Mackenzie, A., McClenaghan, C., Aryal, P., Dong, L., Quigley, A., Grieben, M., Goubin, S., Mukhopadhyay, S., et al. (2015). K2P channel gating mechanisms revealed by structures of TREK-2 and a complex with Prozac. Science 347, 1256–1259. 10.1126/science.1261512.

36. Wilcox, M.R., Nigam, A., Glasgow, N.G., Narangoda, C., Phillips, M.B., Patel, D.S., Mesbahi-Vasey, S., Turcu, A.L., Vazquez, S., Kurnikova, M.G., and Johnson, J.W. (2022). Inhibition of NMDA receptors through a membrane-to-channel path. Nat Commun 13, 4114. 10.1038/s41467-022-31817-z.

37. Kudaibergenova, M., Perissinotti, L.L., and Noskov, S.Y. (2019). Lipid roles in hERG function and interactions with drugs. Neurosci Lett 700, 70–77. 10.1016/j.neulet.2018.05.019.

38. Ahern, C.A., Payandeh, J., Bosmans, F., and Chanda, B. (2016). The hitchhiker’s guide to the voltage-gated sodium channel galaxy. J Gen Physiol 147, 1–24. 10.1085/jgp.201511492.

39. Kim, J.J., and Hibbs, R.E. (2021). Direct Structural Insights into GABA(A) Receptor Pharmacology. Trends Biochem Sci 46, 502–517. 10.1016/j.tibs.2021.01.011.

40. Korner, J., Albani, S., Sudha Bhagavath Eswaran, V., Roehl, A.B., Rossetti, G., and Lampert, A. (2022). Sodium Channels and Local Anesthetics-Old Friends With New Perspectives. Front Pharmacol 13, 837088. 10.3389/fphar.2022.837088.

41. Laskowski, R.A., and Swindells, M.B. (2011). LigPlot+: multiple ligand-protein interaction diagrams for drug discovery. J Chem Inf Model 51, 2778–2786. 10.1021/ci200227u.

42. Elegheert, J., Behiels, E., Bishop, B., Scott, S., Woolley, R.E., Griffiths, S.C., Byrne, E.F.X., Chang, V.T., Stuart, D.I., Jones, E.Y., et al. (2018). Lentiviral transduction of mammalian cells for fast, scalable and high-level production of soluble and membrane proteins. Nat Protoc 13, 2991–3017. 10.1038/s41596-018-0075-9.

43. Mastronarde, D.N. (2005). Automated electron microscope tomography using robust prediction of specimen movements. J Struct Biol 152, 36–51. 10.1016/j.jsb.2005.07.007.

44. Zheng, S.Q., Palovcak, E., Armache, J.P., Verba, K.A., Cheng, Y., and Agard, D.A. (2017). MotionCor2: anisotropic correction of beam-induced motion for improved cryo-electron microscopy. Nat Methods 14, 331–332. 10.1038/nmeth.4193.

45. Rohou, A., and Grigorieff, N. (2015). CTFFIND4: Fast and accurate defocus estimation from electron micrographs. J Struct Biol 192, 216–221. 10.1016/j.jsb.2015.08.008.

46. Wagner, T., Merino, F., Stabrin, M., Moriya, T., Antoni, C., Apelbaum, A., Hagel, P., Sitsel, O., Raisch, T., Prumbaum, D., et al. (2019). SPHIRE-crYOLO is a fast and accurate fully automated particle picker for cryo-EM. Commun Biol 2, 218. 10.1038/s42003-019-0437-z.

47. Scheres, S.H. (2012). RELION: implementation of a Bayesian approach to cryo-EM structure determination. J Struct Biol 180, 519–530. 10.1016/j.jsb.2012.09.006.

48. Punjani, A., Rubinstein, J.L., Fleet, D.J., and Brubaker, M.A. (2017). cryoSPARC: algorithms for rapid unsupervised cryo-EM structure determination. Nat Methods 14, 290–296. 10.1038/nmeth.4169.

49. Morin, A., Eisenbraun, B., Key, J., Sanschagrin, P.C., Timony, M.A., Ottaviano, M., and Sliz, P. (2013). Collaboration gets the most out of software. Elife 2, e01456. 10.7554/eLife.01456.

50. Mirdita, M., Schutze, K., Moriwaki, Y., Heo, L., Ovchinnikov, S., and Steinegger, M. (2022). ColabFold: making protein folding accessible to all. Nat Methods 19, 679–682. 10.1038/s41592-022-01488-1.

51. Liebschner, D., Afonine, P.V., Baker, M.L., Bunkoczi, G., Chen, V.B., Croll, T.I., Hintze, B., Hung, L.W., Jain, S., McCoy, A.J., et al. (2019). Macromolecular structure determination using X-rays, neutrons and electrons: recent developments in Phenix. Acta Crystallogr D Struct Biol 75, 861–877. 10.1107/S2059798319011471.

52. Emsley, P., Lohkamp, B., Scott, W.G., and Cowtan, K. (2010). Features and development of Coot. Acta Crystallogr D Biol Crystallogr 66, 486–501. 10.1107/S0907444910007493.

53. Croll, T.I. (2018). ISOLDE: a physically realistic environment for model building into low-resolution electron-density maps. Acta Crystallogr D Struct Biol 74, 519–530. 10.1107/S2059798318002425.

54. Konagurthu, A.S., Whisstock, J.C., Stuckey, P.J., and Lesk, A.M. (2006). MUSTANG: a multiple structural alignment algorithm. Proteins 64, 559–574. 10.1002/prot.20921.

55. Minh, B.Q., Schmidt, H.A., Chernomor, O., Schrempf, D., Woodhams, M.D., von Haeseler, A., and Lanfear, R. (2020). IQ-TREE 2: New Models and Efficient Methods for Phylogenetic Inference in the Genomic Era. Mol Biol Evol 37, 1530–1534. 10.1093/molbev/msaa015.

56. Le, S.Q., and Gascuel, O. (2008). An improved general amino acid replacement matrix. Mol Biol Evol 25, 1307–1320. 10.1093/molbev/msn067.

57. Ashkenazy, H., Erez, E., Martz, E., Pupko, T., and Ben-Tal, N. (2010). ConSurf 2010: calculating evolutionary conservation in sequence and structure of proteins and nucleic acids. Nucleic Acids Res 38, W529–533. 10.1093/nar/gkq399.

58. Glaser, F., Pupko, T., Paz, I., Bell, R.E., Bechor-Shental, D., Martz, E., and Ben-Tal, N. (2003). ConSurf: identification of functional regions in proteins by surface-mapping of phylogenetic information. Bioinformatics 19, 163–164. 10.1093/bioinformatics/19.1.163.

59. Smart, O.S., Neduvelil, J.G., Wang, X., Wallace, B.A., and Sansom, M.S. (1996). HOLE: a program for the analysis of the pore dimensions of ion channel structural models. J Mol Graph 14, 354–360, 376. 10.1016/s0263-7855(97)00009-x.

60. Trott, O., and Olson, A.J. (2010). AutoDock Vina: improving the speed and accuracy of docking with a new scoring function, efficient optimization, and multithreading. J Comput Chem 31, 455–461. 10.1002/jcc.21334.

61. Eberhardt, J., Santos-Martins, D., Tillack, A.F., and Forli, S. (2021). AutoDock Vina 1.2.0: New Docking Methods, Expanded Force Field, and Python Bindings. J Chem Inf Model 61, 3891–3898. 10.1021/acs.jcim.1c00203.

