## Supplemental Figures for "Structure of an insect gustatory receptor"

### Supplementary Figures

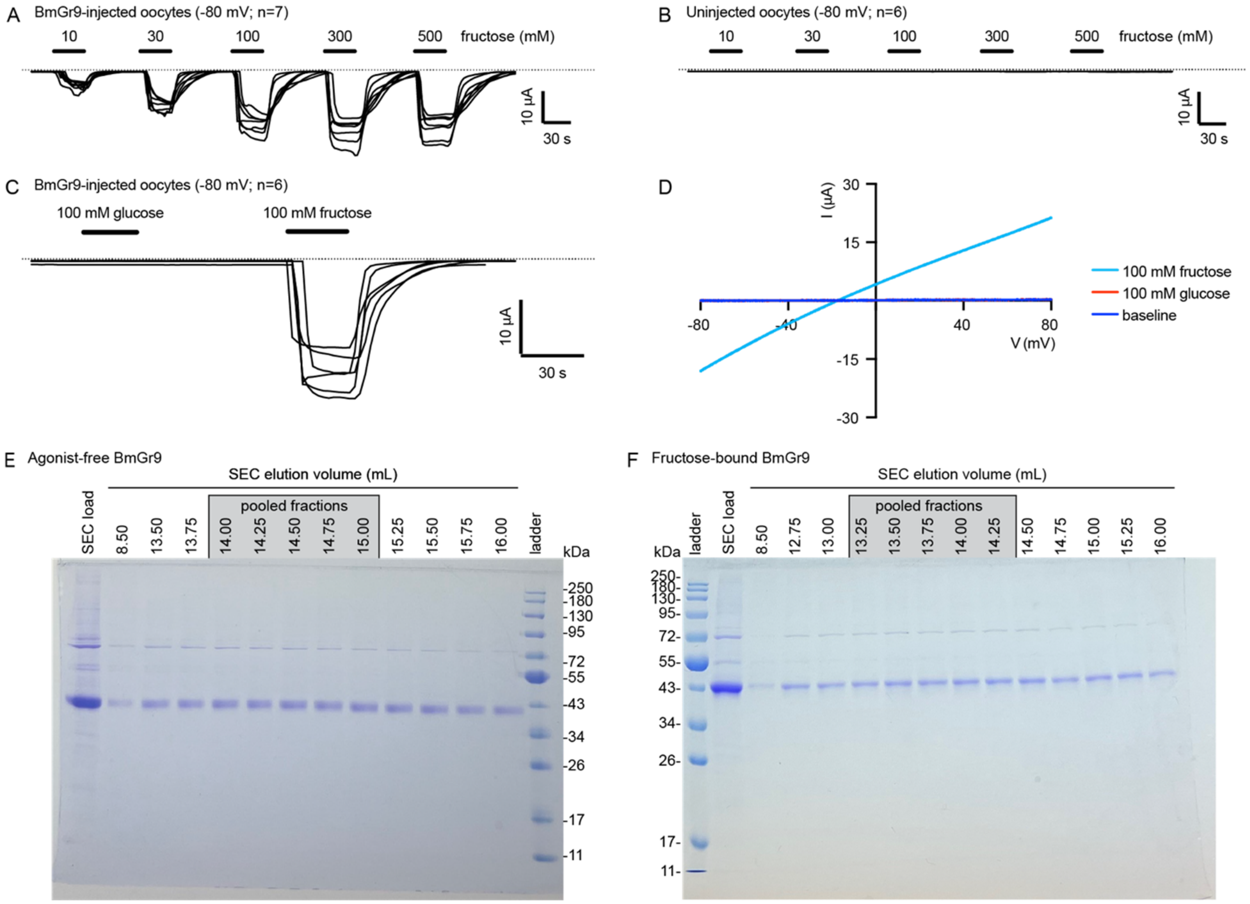


**Figure S1. Functional analysis and purification of Twin-Strep-tagged BmGr9**

(A-C) BmGr9-expressing oocytes respond specifically to fructose. Individual traces show current at -80 mV in different oocytes. (A) BmGr9-expressing oocytes (n=7 oocytes). (B) Uninjected control oocytes (n=6). (C) BmGr9-expressing oocytes (n=6).

(D) Current-voltage relationships of BmGr9-expressing oocytes.

(E-F) Coomassie-stained SDS-PAGE gels of the size exclusion chromatography (SEC; Superose 6 10/30) load and elution fractions of agonist-free BmGr9 (E) and fructose-bound BmGr9 (F). The corresponding elution volumes are above each lane, and the fractions pooled and concentrated for cryo-EM are indicated. The SEC load fractions were diluted 1:10.

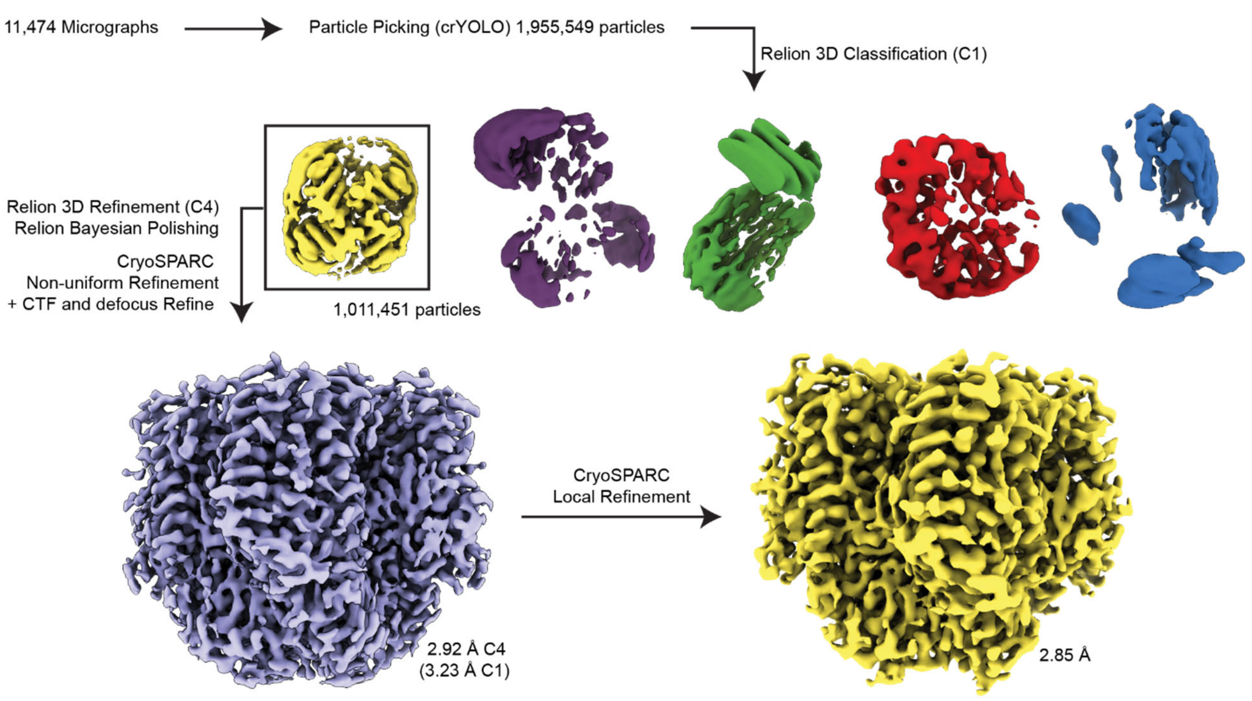


**Figure S2. Cryo-EM processing procedure for the agonist-free BmGr9 structure**

Processing scheme for classification and refinement of the agonist-free BmGr9 map. The locally filtered map used for the final reconstruction is represented with dust hidden for clarity.

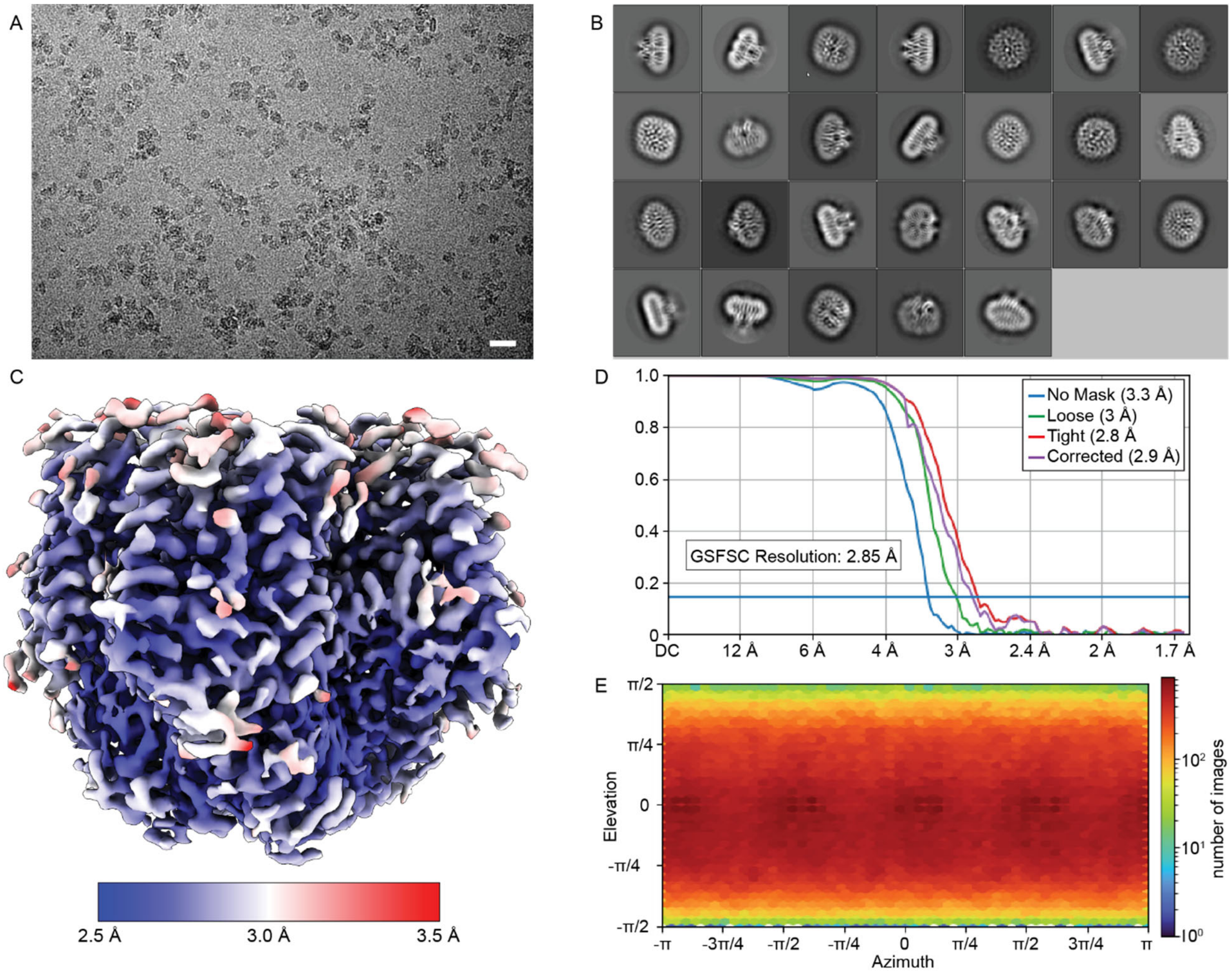


**Figure S3. Cryo-EM data analysis for the agonist-free BmGr9 structure**

(A) Representative micrograph of BmGr9 embedded in vitreous ice (scale bar = 250 Å), low pass filtered for clarity.

(B) Selected 2D class averages of BmGr9.

(C) Reconstruction of BmGr9 filtered and colored by local resolution.

(D) Gold-standard Fourier shell correlation (FSC) curves from cryoSPARC.

(E) Viewing direction distribution plot.

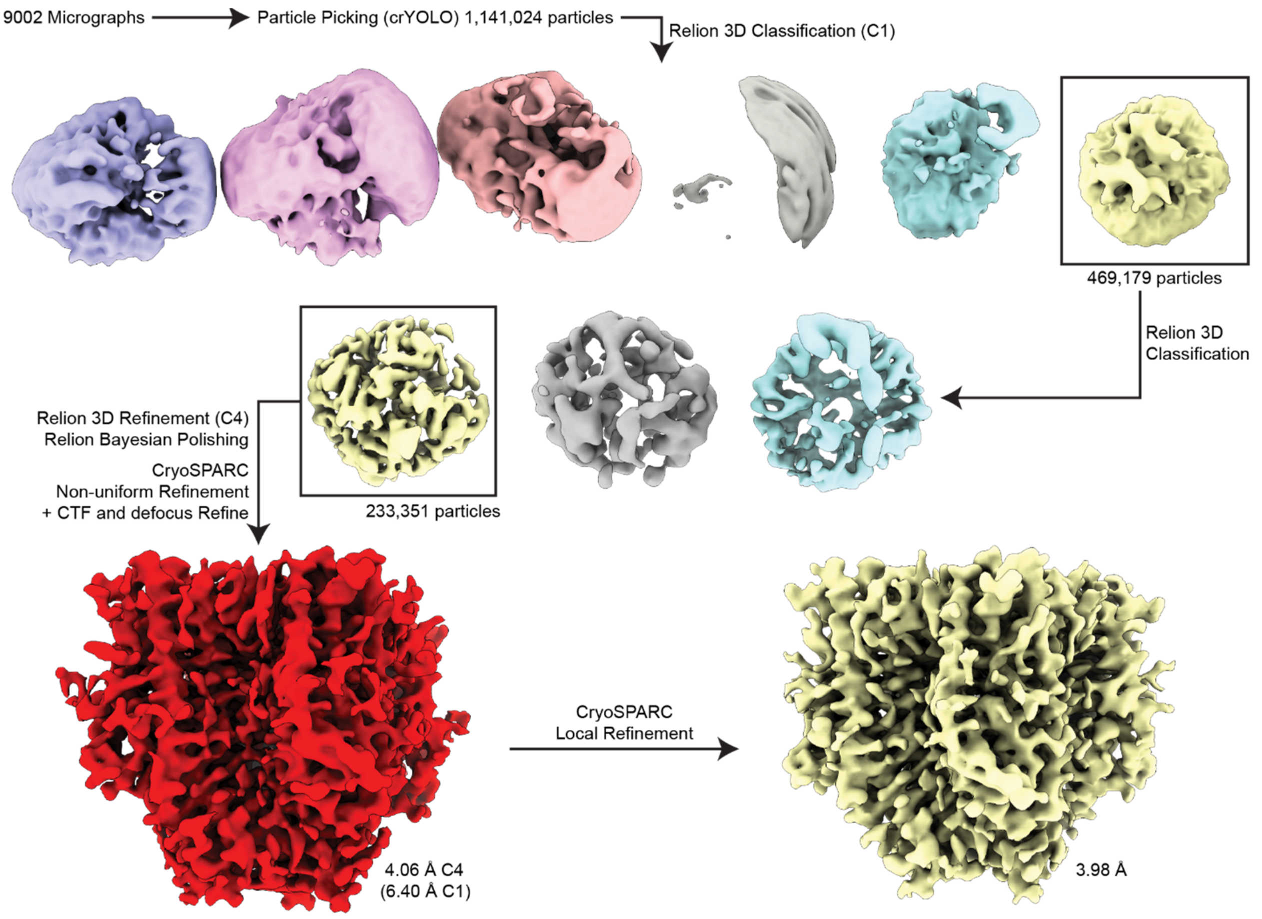


**Figure S4. Cryo-EM processing procedure for the fructose-bound BmGr9 structure**

Processing scheme for classification and refinement of the fructose-bound BmGr9 map. The locally filtered map used for the final reconstruction is represented with dust hidden for clarity.

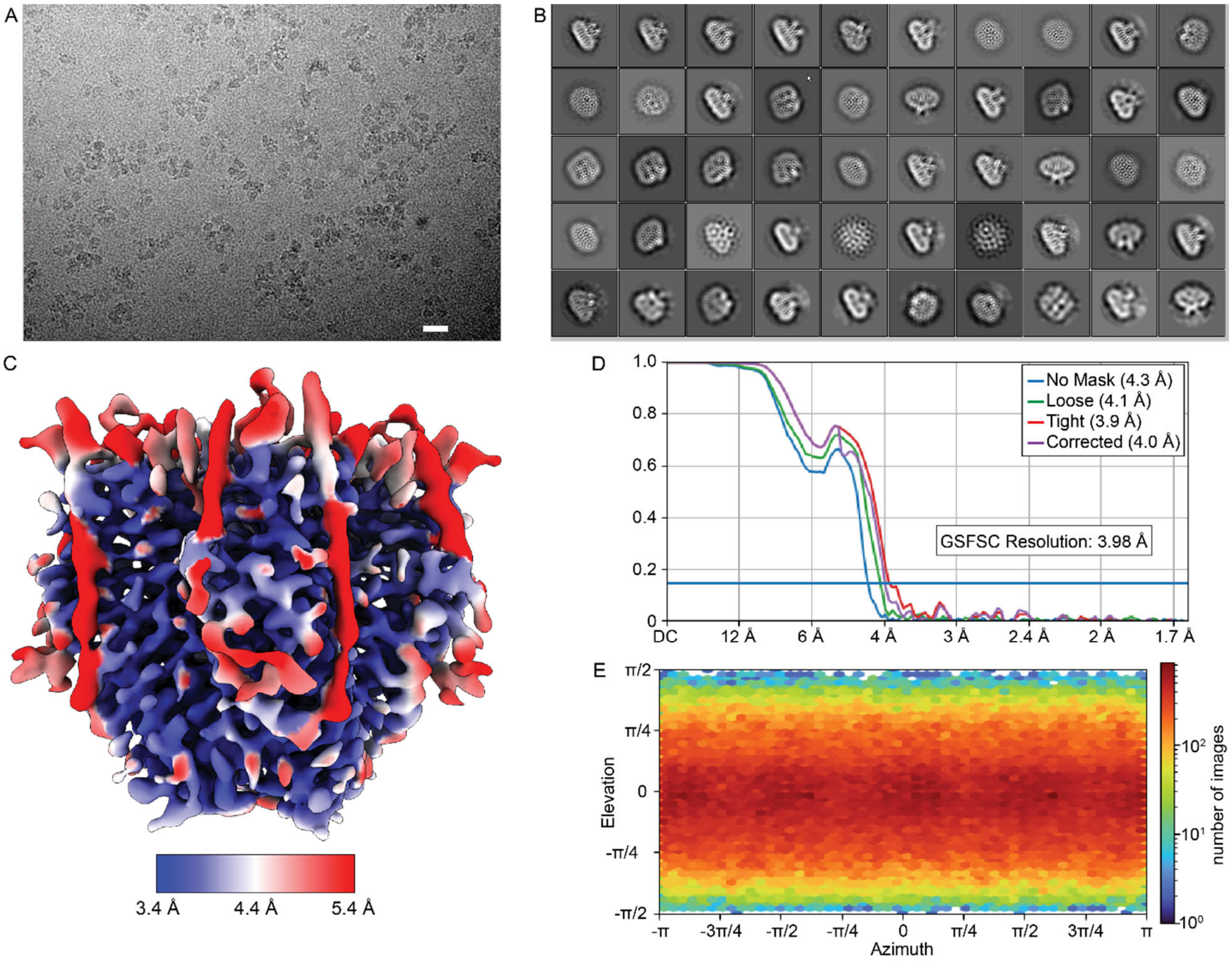


**Figure S5. Cryo-EM data analysis for the fructose-bound BmGr9 structure**

**(**A) Representative micrograph of BmGr9 in the presence of fructose embedded in vitreous ice (scale bar = 250 Å), low pass filtered for clarity.

(B) Selected 2D class averages of fructose-bound BmGr9.

(C) Reconstruction of fructose-bound BmGr9 filtered and colored by local resolution.

(D) Gold-standard Fourier shell correlation (FSC) curves from cryoSPARC.

(E) Viewing direction distribution plot.

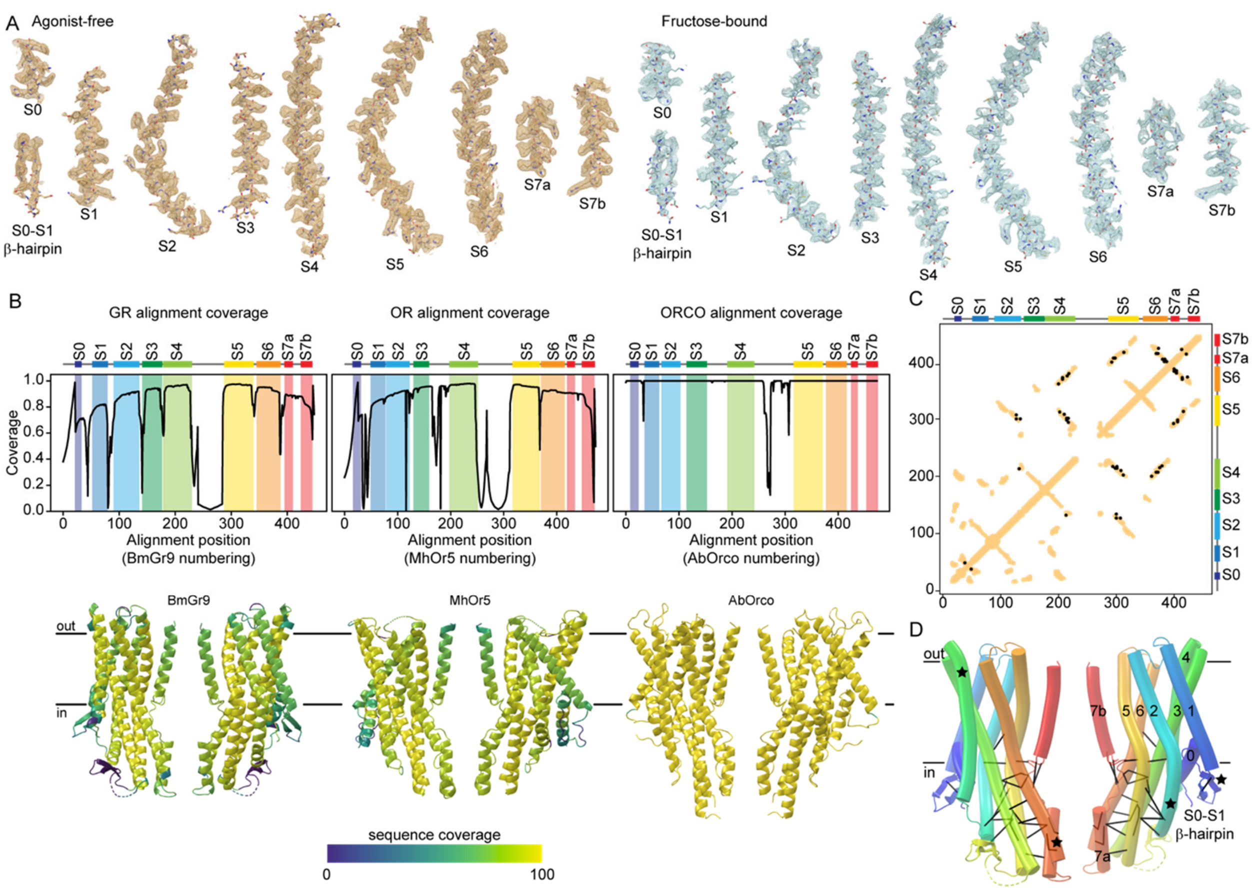

**Figure S6. Structural and sequence data supporting secondary structure and topology analyses**

(A) Cryo-EM densities of individual helices for the agonist-free (left, orange) and fructose-bound (right, blue) BmGr9 structures. Protein is shown in stick representation with density contoured to 3.5σ and 4.5σ for the agonist-free and fructose-bound maps, respectively. The helices (and S0-S1 β-hairpin) are labeled below each helix from N to C terminus.

(B) Sequence coverage for the three sequence alignments mapped onto the sequence (top) and structure (bottom) of the representative member, as follows: structure-driven alignment of 1895 GR sequences (includes 41 OR sequences) mapped to BmGr9; structure-driven alignment of 3885 OR sequences mapped to MhOr5; and previously published alignment of 176 ORCO sequences[24] mapped to AbOrco.

(C) Sequence covariation analysis using the structure-driven alignment of 1854 GR sequences mapped to BmGr9. Intrasubunit structural contacts in BmGr9 are in orange (8 Å cutoff). The 28 evolutionarily coupled residue pairs above the 90% confidence threshold are indicated by black dots.

(D) The 28 evolutionarily coupled residues pairs mapped onto the BmGr9 structure as black Cα-to-Cα bonds. Two opposing subunits are shown in transparent cartoon representation colored in blue-to-red rainbow from N to C terminus. Nearly all top coupled pairs are in the intracellular anchor domain. Noteworthy structural features highlighted in Figure 2A are marked with black stars.

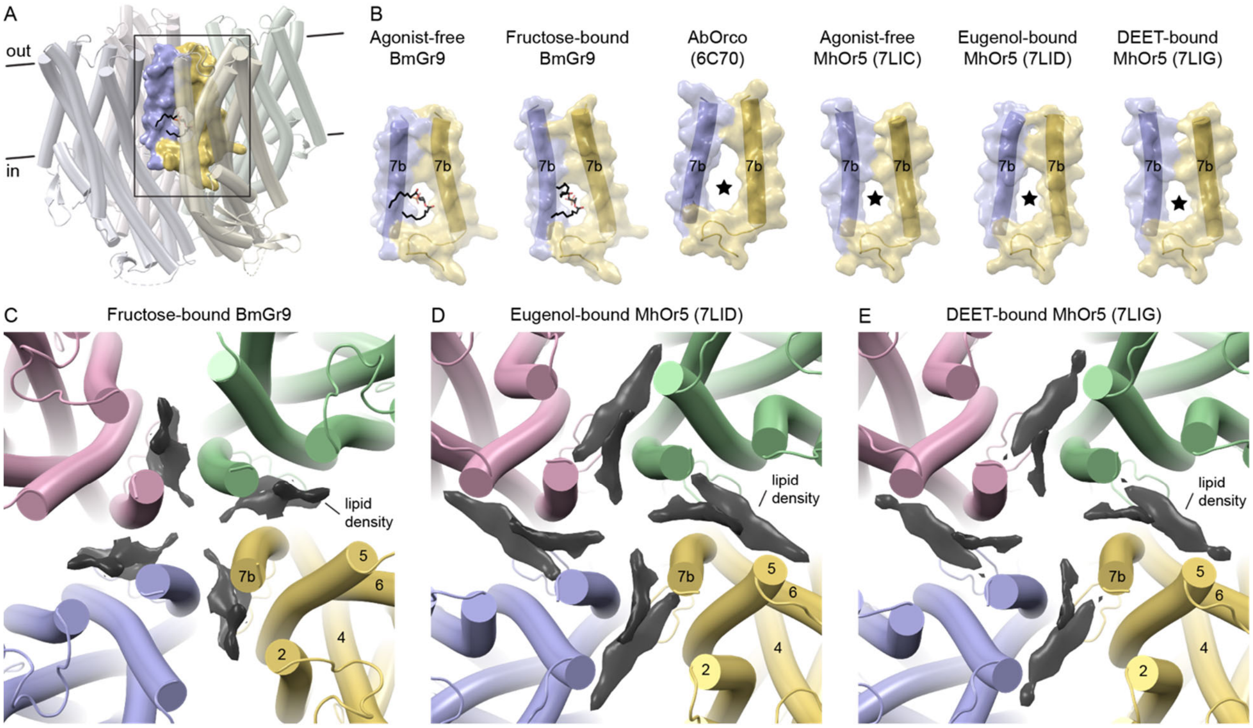


**Figure S7. Density for a lipid penetrating the ion pore is observed in other GR and OR structures**

(A) Structure of agonist-free BmGr9 illustrating the position of the transmembrane fenestration between the blue and yellow subunits with the bound pore-penetrating lipid (black sticks). The boxed region is shown in B for this structure and other available structures of GRs and ORs.

(B) All GR and OR structures have fenestrations between pore helices in the membrane plane. For each available GR or OR structure, the transparent surface and cartoon representation of helix S7b from two adjacent subunits and the loop preceding helix S7b for the yellow subunit are illustrated. For the two BmGr9 structures, the modeled pore-penetrating lipid is shown in black sticks; for the other structures, the fenestration is marked by a black star.

(C-E) Transmembrane cross-sections of BrGr9 and MhOr5 structures viewed from the extracellular side of the membrane. Surfaces shown in black are densities in the respective cryo-EM map corresponding to potential pore-penetrating lipids. The fructose-bound BmGr9 structure and map contoured at 3.5σ are shown in (C); the eugenol-bound MhOr5 structure (PDB ID: 7LID) and map (EMDB ID: 23374) contoured at 7.5σ in (D); and the DEET-bound MhOr5 (PDB ID: 7LIG) and map (EMDB ID: 23375) contoured at 7.5σ in (E). Visible helices of the yellow subunit are labeled.

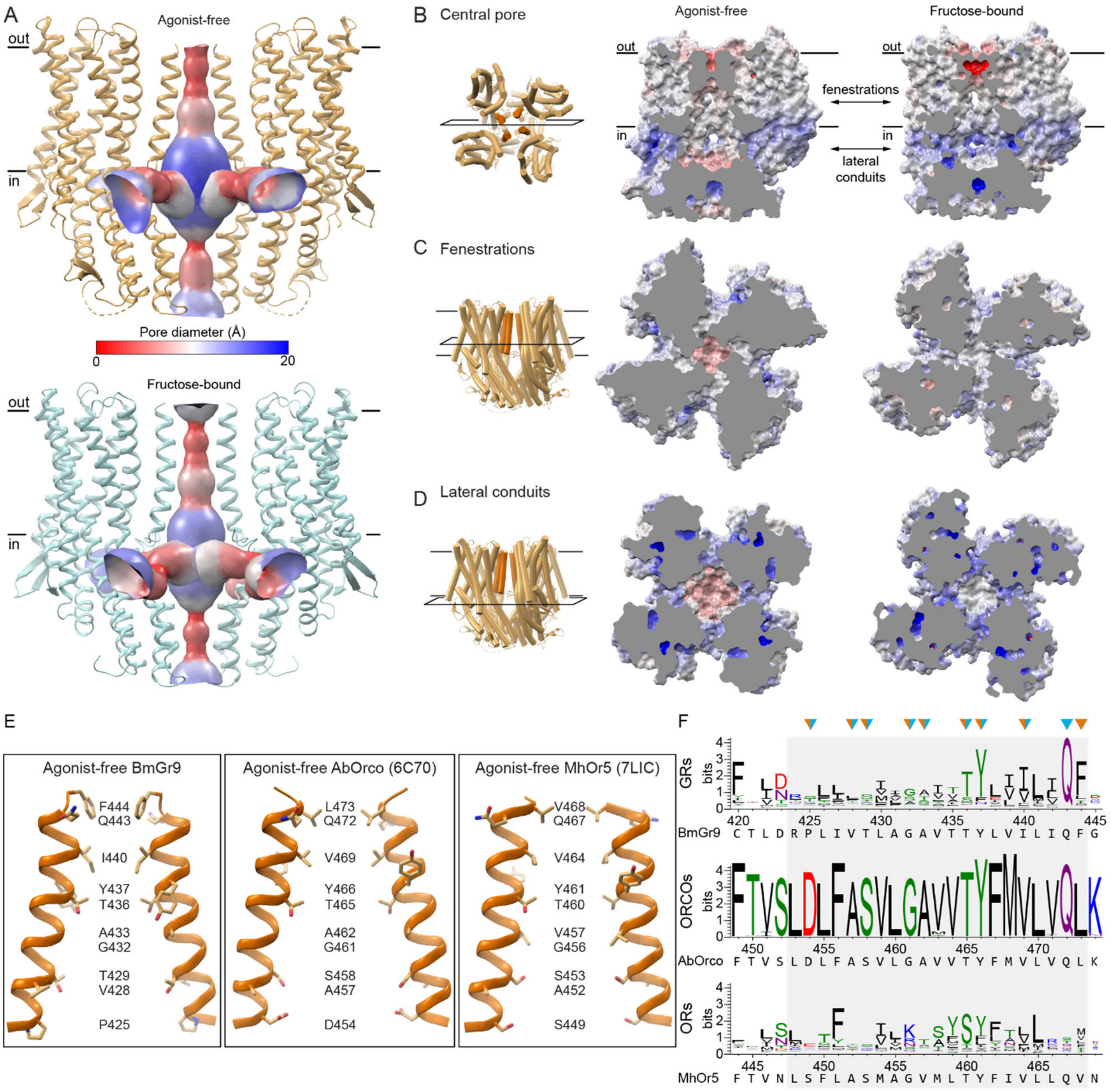


**Figure S8. Comparisons of pore between BmGr9 and AbOrco and MhOr5**

(A) The quadrivial pore of BmGr9 in the agonist-free (top) and fructose-bound structures illustrated as a surface colored according to its diameter. The cartoon representation of two opposing protein subunits is included, and black lines make the membrane boundaries.

(B-D) Electrostatics surface representation of BmGr9 in the agonist-free (left) and fructose-bound (right) structures, in three cross-sections as indicated by the cartoon representations and planes on the left: (B) Vertical cross-sections through the ion pore. Black lines mark the membrane boundaries, and arrows point to the hydrophobic lateral fenestrations filled with the pore-penetrating lipids and the lateral conduits of the ion pore. (C) Horizontal cross-sections through the hydrophobic fenestrations. (D) Horizontal cross-sections through the lateral conduits. The electrostatics potentials were calculated using APBS in PyMOL and colored as a range from -20 kcal/mol (red) to 20 kcal/mol (blue).

(E) Comparison of the central ion pore of the agonist-free structures of BmGr9 (left), AbOrco (middle; PDB ID: 6C70), and MhOr5 (right; PDB ID: 7LIC). Helix S7b from two opposing protein subunits are shown in cartoon representation, and the sidechains of residues that line the pore walls are shown in sticks and labeled.

(F) Sequence logos of the helix S7b positions (grey box) of alignments of 1854 insect GR sequences, 176 ORCO sequences, and 3885 insect OR sequences, with the reference sequences of BmGr9, AbOrco, and MhOr5, respectively, indicated below the logo. Residues lining the central pore are marked with arrowheads colored orange (agonist-free structure) or blue (fructose-bound structure) or both colors for sidechains that are pore-lining in both structures.

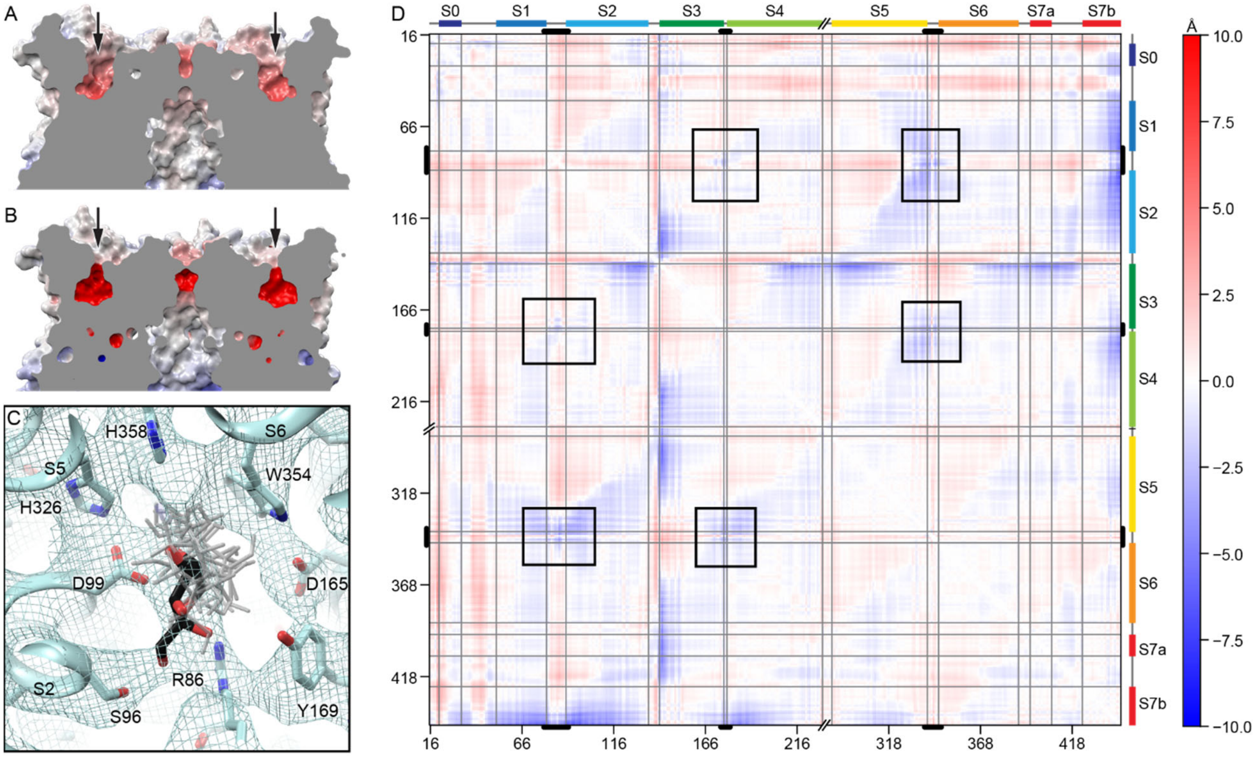


**Figure S9. Conformational changes in and fructose docking into the BmGr9 ligand-binding pocket**

(A-B) Electrostatics surface representation of agonist-free (A) and fructose-bound (B) BmGr9, sliced through the two ligand-binding pockets of opposing subunits in the tetramer. The two pockets are indicated with black arrows. The electrostatics potentials were calculated using APBS in PyMOL and colored as a range from -20 kcal/mol (red) to 20 kcal/mol (blue).

(C) Cryo-EM density for the fructose-bound map contoured at 4.5σ around the fructose-binding site, with nearby sidechains shown as sticks. The five lowest-energy poses of docked β-D-fructopyranose are illustrated in grey sticks, and the β-D-fructopyranose pose after real-space refinement of the lowest-energy pose against the cryo-EM density is shown in black sticks.

(D) Distance difference matrix of Cα-to-Cα distances for agonist-free versus fructose-bound BmGr9. The breaks in the axes mark missing residues 230-283. Thick black lines on the four axes mark the extracellular loops, and grey lines mark the helix boundaries. Black boxes mark the relative distances between the three pocket-forming regions of BmGr9, highlighting that the extracellular regions generally move closer together upon fructose binding.

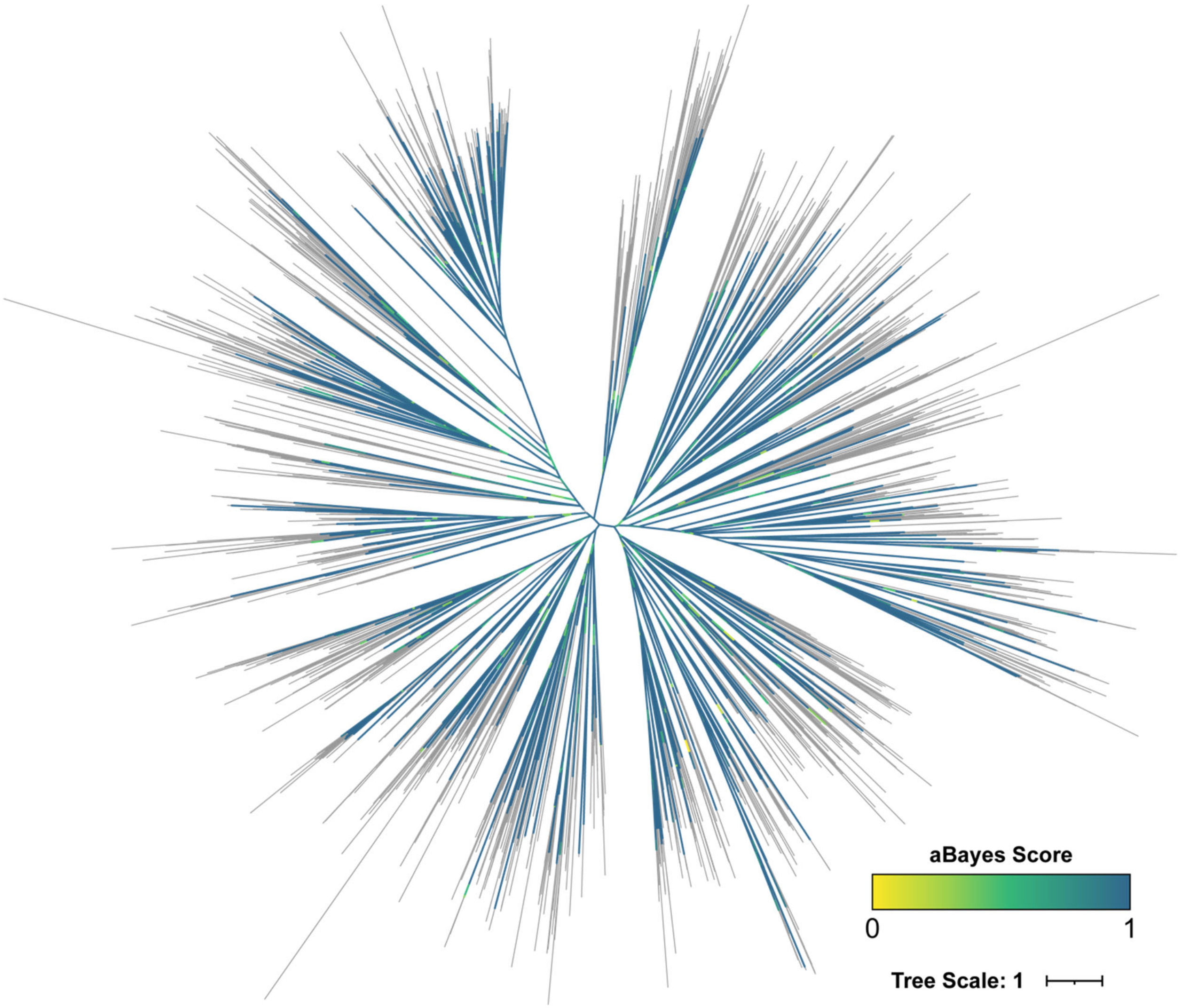


**Figure S10. Posterior probability branch supports for the maximum likelihood phylogenetic tree of insect GR sequences**

A representation of the phylogenetic tree in Figure 6A showing aBayes support values for each branch. Terminal branches are shown in grey.

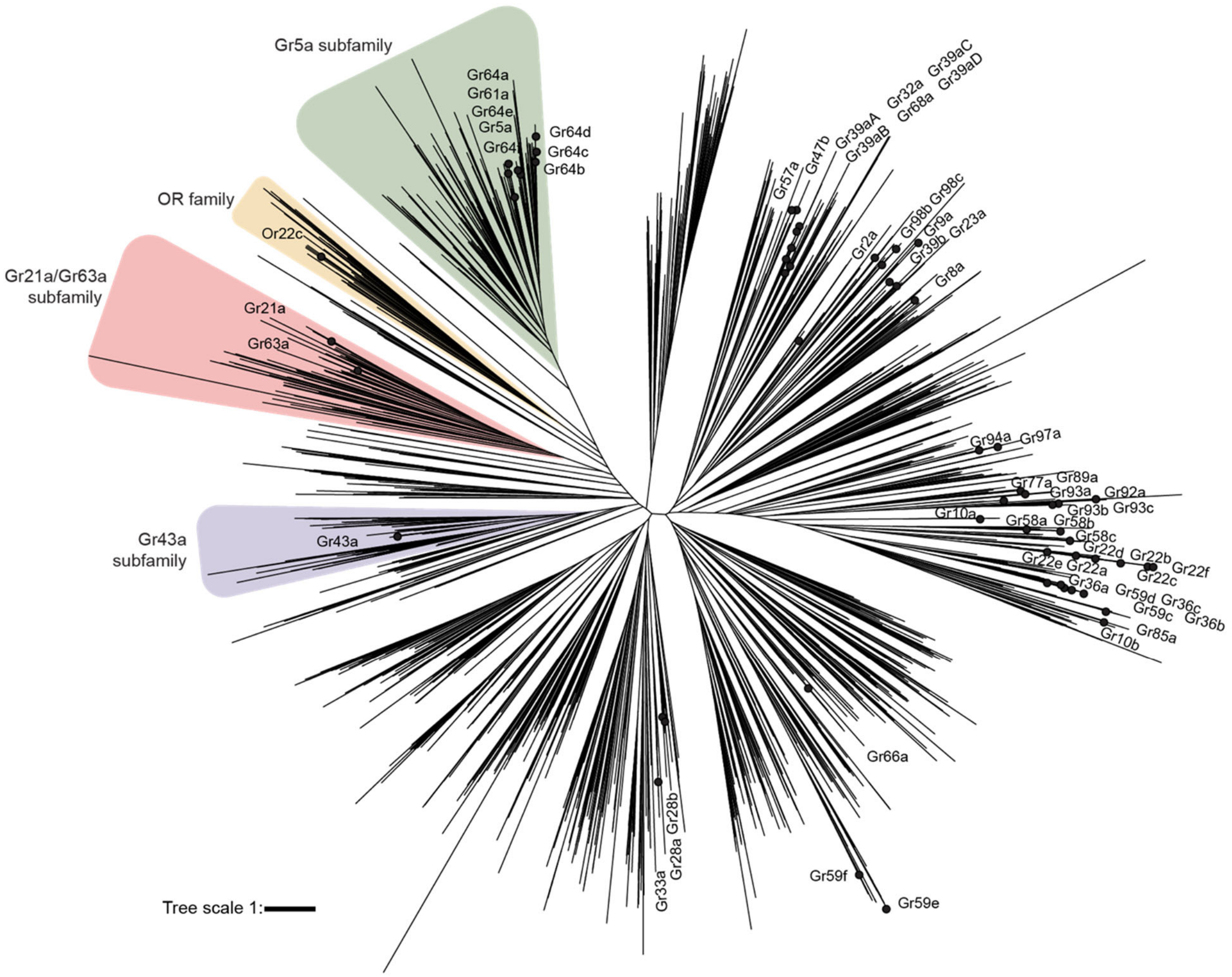


**Figure S11. Annotated phylogenetic tree of insect GR sequences**

Maximum likelihood phylogenetic tree of 1895 aligned insect GR sequences. Black dots indicate positions of *D. melanogaster* GRs on the tree. The alignment included 41 OR sequences, marking the branch point of the OR family within the more ancestral GR sequences (orange). The three subfamilies analyzed in this work are highlighted: Gr5a subfamily in green, Gr63a subfamily in red, and Gr43a subfamily in purple.

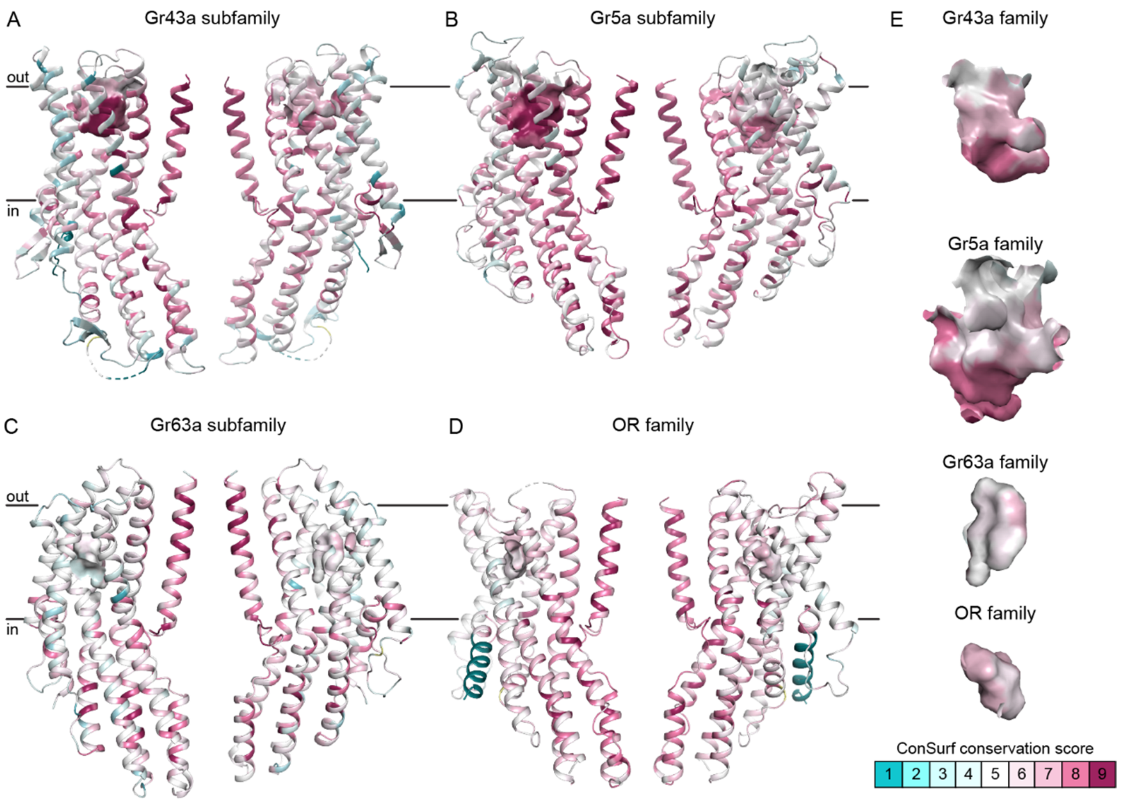


**Figure S12. Side views of predicted ligand-binding pockets in representative GR subfamily members**

(A-E) Cartoon representation of two opposing subunits from representative subfamily members viewed from the membrane plane, with surface representation of the atoms forming the predicted ligand-binding pockets to illustrate the relative location and size of the binding pockets. The cartoons and surfaces are colored by ConSurf conservation score based on the sequence alignment of the corresponding GR subfamily. As in Figure 6, the following subunits and corresponding protein (sub)families are illustrated: (A) Gr43a subfamily (74 sequences) on the agonist-free BmGr9 structure; (B) Gr5a subfamily (251 sequences) on the AlphaFold2 model of Gr5a (UniProt ID: Q9W497); (C) Gr63a subfamily (107 sequences) on the AlphaFold2 model of Gr63a (UniProt ID: Q9VZL7); and (D) the OR family (3885 sequences) on the agonist-free MhOr5 structure (PDB ID: 7LIC). For the AlphaFold2 models, tetramers were built for visualization purposes by superposition of four subunit models on the BmGr9 tetramer.

(E) Zoomed-in views of the predicted ligand-binding pockets from the right-hand subunit of panels A-D to highlight relative size of the pockets across (sub)families. Some stray surfaces not corresponding to the pocket volumes were removed for clarity.

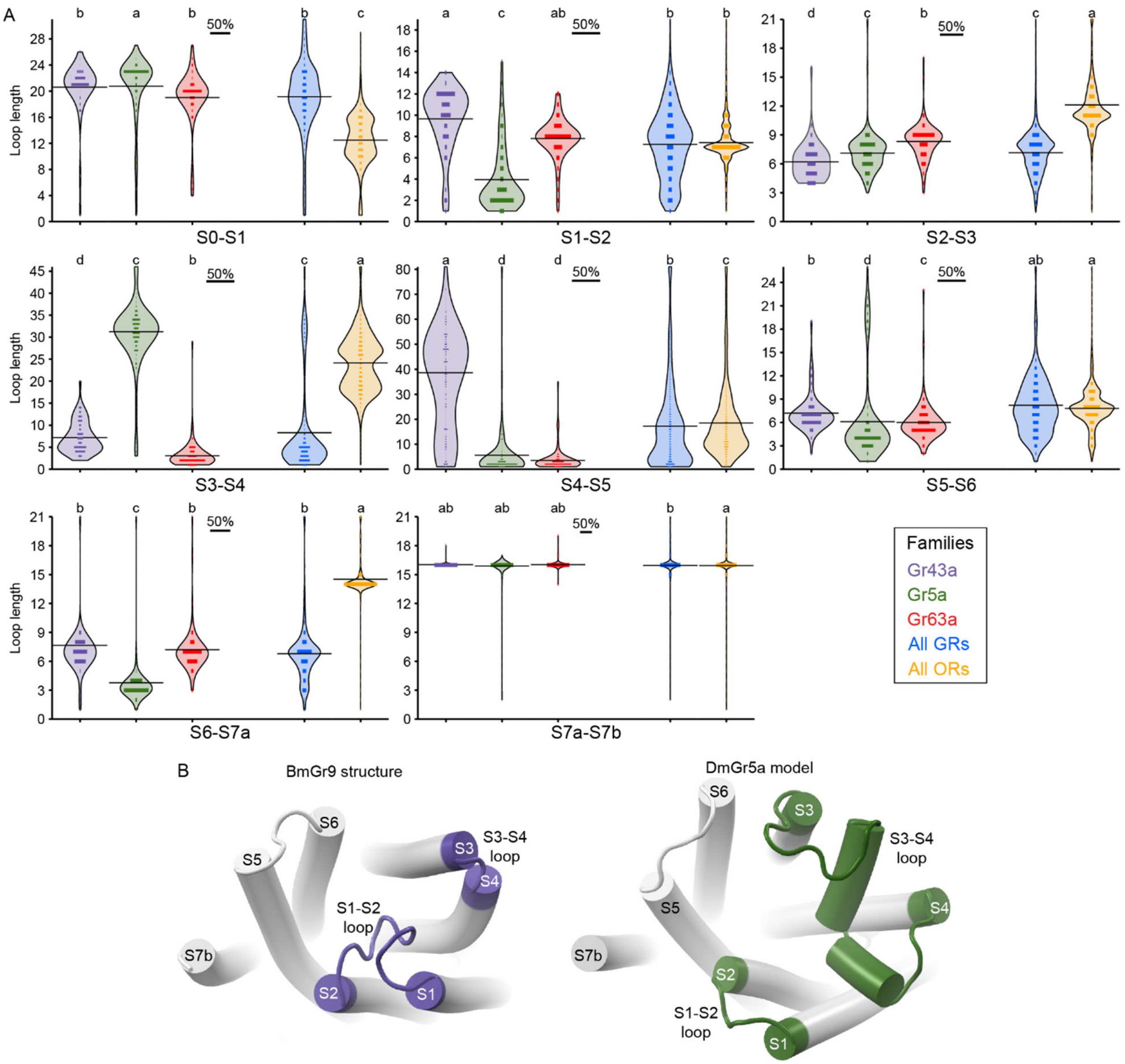


**Figure S13. Distribution of loop lengths across GR subfamilies, the GR family and the OR family**

(A) Violin plots depict the distribution of lengths for each loop connecting two adjacent helices as indicated in each plot. Each plot shows the loop length (in number of residues) on the y axis and receptor (sub)family types on the x axis. The length of the lines inside the violins is proportional to the fraction of sequences in each (sub)family with the given loop lengths, with the 50% scale bars indicated at the top of each plot. Each plot shows five different violins: Gr43a subfamily (violet), Gr5a subfamily (green), Gr63a subfamily (red), all GRs (blue), and all ORs (orange). Distinct letters above each violin denote statistically distinct classes, listed alphabetically from highest median value to lowest such that adjacent categories bear adjacent letters based on a Steel-Dwass test, p<0.01.

(B) Top views of a single subunit from the agonist-free BmGr9 structure (left) and the DmGr5a AlphaFold2 model (right) showing relative differences in length and structure of their S1-S2 and S3-S4 loops.
